# TreeGOER: a database with globally observed environmental ranges for 48,129 tree species

**DOI:** 10.1101/2023.05.15.540790

**Authors:** Roeland Kindt

**Affiliations:** Trees and forest genetic resources and biodiversity, World Agroforestry, CIFOR-ICRAF

## Abstract

The BIOCLIM algorithm provides a straightforward method to estimate the effects of climate change on the distribution of species. Estimating the core ranges of species from 5% and 95% quantiles of bioclimatic variables, the algorithm remains widely used even when more sophisticated methods of species distribution modelling have become popular. Where sufficient representative observations are available, I expect that BIOCLIM correctly identifies locations that would not be suitable in a future climate. To accommodate climate change investigations based on BIOCLIM for 48,129 tree species (a substantial subset of known tree species), I developed the `TreeGOER` (Tree Globally Observed Environmental Ranges) database, providing information on environmental ranges for 38 bioclimatic, 8 soil and 3 topographic variables. The database can be accessed from: https://doi.org/10.5281/zenodo.7922928. Statistics that include 5% and 95% quantiles were estimated for a cleaned and taxonomically standardized occurrence data set with different methods of outlier detection, with estimates for roughly 45% of species being based on 20 or more observation records. Inferred core bioclimatic ranges of species along global temperature and moisture index gradients and across continents follow the known global distribution of tree diversity such as its highest levels in moist tropical forests and the ‘odd man out’ pattern of lower levels in Africa. To demonstrate how global analyses for large numbers of tree species can easily be done in `R` with `TreeGOER`, here I present two case studies. The first case study investigated latitudinal trends of tree vulnerability and compared these with previous results obtained for urban trees. The second case study focused on tropical areas, compared trends in different longitudinal zones and investigated patterns for the moisture index. `TreeGOER` is expected to benefit researchers conducting biogeographical and climate change research for a wide range of tree species at a variety of spatial and temporal scales.

## 1 Introduction

Trees are of immense importance to ecological systems, the global economy and to human livelihoods and wellbeing (Di Sacco et al., 2021; Rivers et al., 2022). At least a quarter of known tree species have been documented to be useful (Kindt et al., 2023), whereby they provide the matrix of many terrestrial ecosystems, define agroforestry systems and play key roles in climate regulation through carbon and water cycling (Keppel et al., 2021; van Noordwijk et al., 2021). Given their importance and the ensuing climate change crisis (Ripple et al., 2017; Lyon et al., 2021), it has become essential to anticipate the effects of climate change on their future suitability. Estimating the future suitability of tree species is not only relevant to devise adaptation strategies for trees in forests and agroforestry systems (Meybeck et al., 2021), but also provides insights in mitigation pathways by linking assemblages of future-suitable species with their carbon sequestration potentials (Jucker et al., 2022; Rius et al., 2023; Duguma et al., 2023). It is equally important that massive ecological restoration initiatives such as the UN Decade on Ecosystem Restoration or the Bonn Challenge (Chapman et al., 2020; Höhl et al., 2020; van Noordwijk et al., 2020) factor in climate change effects – this has been explicitly included in the *International Principles and Standards for the Practice of Ecological Restoration* via Principle 3 that *“Ecological Restoration Practice Is Informed by Native Reference Ecosystems, while Considering Environmental Change”* (Gann et al., 2019).

To estimate the suitability of tree species in future climates, species distribution models (SDMs) need to be calibrated, and to calibrate these models, observations are required that represent the range of environmental conditions under which a species can occur. Booth (2018) has argued that for most tree species, the only option to learn about climate change impacts are correlative SDMs such as models described by Guisan et al. (2017). For each tree species, presence observations are therefore required which ideally should characterize the full range of environmental conditions where a species can occur.

GlobalTreeSearch (GTS) was the first global database aimed at listing all known tree species and is updated regularly (Beech et al., 2017; https://tools.bgci.org/global_tree_search.php). GTS uses the tree definition of IUCN’s Global Tree Specialist Group of *“a woody plant with usually a single stem growing to a height of at least two metres, or if multi-stemmed, then at least one vertical stem five centimetres in diameter at breast height”*. GTS excludes hybrid species, cycads, tree ferns and tree-like Poaceae, Bromeliaceae and Musaceae species. Updated regularly, the most recent downloadable version listed 57,958 species (version 1.6 of April 2022). Searching for available occurrence records for all known tree species, Serra-Diaz et al. (2017) found 49,206 tree species with available records from which they retained 15,140 species with at least 20 records after data cleaning. Keppel et al. (2021; see their Appendix S3) compiled a global standardized list of 58,044 tree species informed partially by GTS version 1.4. Their analysis of georeferenced records available in GBIF, the largest database of occurrence data available among those they analyzed, showed that 48,970 species (84.4%) had at least one record.

Achieving acceptable SDM calibrations requires a minimum number of observation records, with this number of records widely debated. Results obtained from Wisz et al. (2008) suggest that a minimum of 30 records are required in combination with superior modelling algorithms to achieve acceptable SDM calibration results. Van Proosdij et al. (2016) in an African study documented lower limits that ranged from 14 for narrow-ranged to 25 for widespread species. Varela et al. (2014) suggest that for optimal performance of non-filtered data, 50 observations may be required but that environmental filtering results in smaller required sample sizes (but obviously filtering requires a larger initial sample size). A substantially higher number of records is suggested by Feeley and Silman (2011) as a consequence of the spatiotemporal aggregation of collections, with their observation that time-sequenced collections of 75-100 occurrences was on average equivalent to 25 randomly subsampled occurrences. Santini et al. (2021) when assessing the reliability of SDM in climate change research made the argument to use large sample sizes (tentatively 200 – 500 points) because of uncertainties for most species about ecologically-meaningful predictor variables, biases in sampling and questions whether a species was in equilibrium with its environment.

Although studies such as Elith et al. (2006) and Wisz et al. (2008) found that more sophisticated SDM algorithms such as MAXENT or boosted regression trees outperformed BIOCLIM, this pioneering method remains widely used, especially through the concept of bioclimatic variables that it developed (Booth, 2018). BIOCLIM also remains to be used within the context of ensemble modelling frameworks (Marmion et al., 2008; Kindt, 2018; Brummit et al., 2020). One of its utilities comes from mapping areas where different algorithms reach consensus that a species would be suitable, thus showing areas with which the easily-understood BIOCLIM agrees. A recent example of using a BIOCLIM algorithm to infer the effects of climate change is the global study of urban tree vulnerability by Esperon-Rodriguez et al (2022); this study will be discussed in further detail among the case studies. The Climate Assessment Tool of Botanic Gardens Conservation International that was launched near the end of 2022 (https://www.bgci.org/resources/bgci-hosted-data-tools/climate-assessment-tool/) also uses a version of the BIOCLIM algorithm, but modified the system to identify locations near the edge of the known distribution through 1% - 10% and 90% - 99% percentile ranges, whereas the default BIOCLIM algorithm uses 5%- 95% quantile ranges to identify the core bioclimatic domain, and minimum – 5% and 95% - maximum ranges to define marginal bioclimatic domains (e.g., Lindenmayer et al., 1996).

Hijmans and Graham (2006) did not recommend using BIOCLIM for climate change investigations and instead endorsed more complex algorithms such as generalized additive models (GAM) and maximum entropy models (MAXENT). However, their arguments (as in their discussion of their Figure 3) combined issues of sample sizes and representative sampling besides methodological differences between algorithms. They stated that *“Bioclim can be used as a conservative approach, for example, in the context of reserve planning. It will likely underestimate future ranges, but there is a high probability that areas identified as suitable for a species will be correctly identified”*. My opinion is that BIOCLIM may instead overestimate future ranges compared to other algorithms (such as GAM and MAXENT). I illustrate my argument via Figure 1 where the environmental niche of a species has an ellipsoid shape as used in previous theoretical discussions as by Hijmans and Graham (2006), Etherington (2019) or Erickson and Smith (2023). The same arguments can be made, however, for niches with convex or concave hull shapes, as used for example in climate change studies by Pironon et al. (2019) or van Zonneveld et al. (2023). With a large enough sample size where more complex model calibrations can approximate the true ellipsoid niches well, the BIOCLIM algorithm tends to overestimate suitable conditions in the zones labelled as ‘B1’ in Figure 1. If one further accepts the additional condition for more complex approaches, that for each individual explanatory variable the conditions at the tails of the distribution should not be predicted to be suitable (put alternatively, accepting the ecological justification of the BIOCLIM algorithm and deciding not to include marginal bioclimatic conditions), then zones labelled as ‘B2’ in Figure 1 would be modelled as not being suitable regardless of the SDM. The overprediction of suitable conditions in the ‘B1’ zones by BIOCLIM becomes more prominent in situations where environmental variables are more strongly correlated (Figure 1a), whereas where variables are correlated less most of the suitable area predicted by BIOCLIM corresponds to the ‘A’ zone (Figure 1b).

**FIGURE 1.**
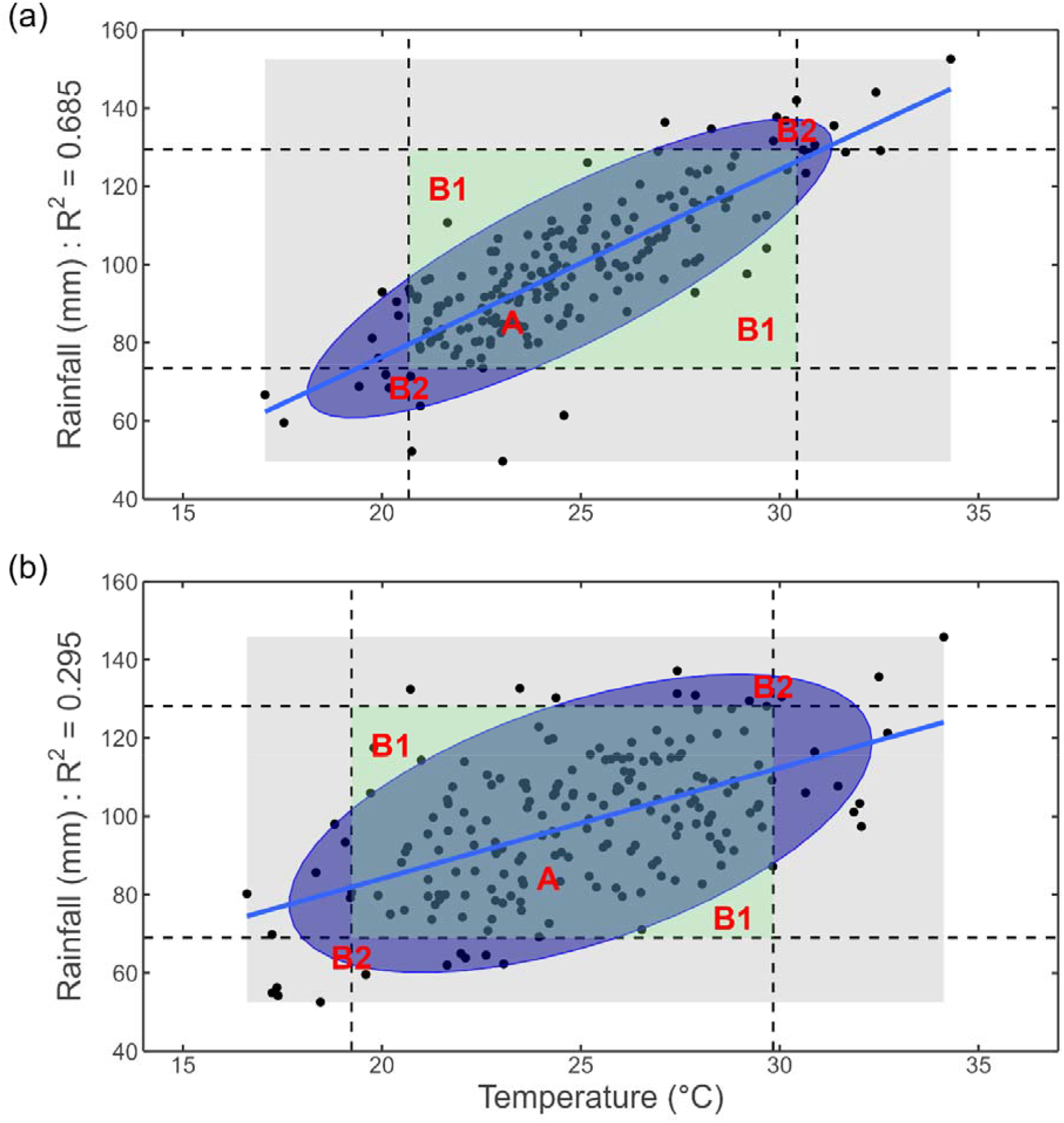
Comparison of environmental niches estimated by BIOCLIM and by an ellipsoid-fitting model where the species’ fundamental niche is elliptically-shaped following a multivariate normal distribution (a) with relatively high correlation between explanatory variables and (b) with relatively low correlation. The distribution of observation points for each subfigure was generated using function `MASS::mvrnorm` similar to `R` scripts used by Etherington (2019) but with different covariance matrices and 200 observations in each simulation. Dashed lines correspond to 5% and 95% quantile limits and delineate the core bioclimatic niche that BIOCLIM identifies. The ellipsoid niche was estimated via `ggplot2::stat_ellipse` with `level` set at 0.90. ‘A’ indicates a zone where BIOCLIM and the ellipsoid-fitting model agree that the species is suitable, ‘B1’ a zone where only BIOCLIM predicts that the species is suitable, and ‘B2’ a zone where only the ellipsoid-fitting model predicts the species is suitable.

A general practice when calibrating species distribution models is to select explanatory variables that are correlated less with methods such as the Variance Inflation Factor analysis (Ranjitkar et al., 2014; de Sousa et al., 2019; Fremout et al., 2020). The justification for these selections come from modelling complications such as overfitting and required sample sizes that multiply with the number of variables used (Erickson and Smith 2023). A shortcoming of selecting a subset of less-correlated variables, however, is that correlations may be different in future climates and that therefore different subsets of variables could generate different future projections (Braunish et al., 2013). My observations from Figure 1 seem to point in a similar direction that using a larger set of more-correlated variables provides an advantage by allowing more complex algorithms to correctly identify the ‘B1’ zones that BIOCLIM fails to identify as unsuitable.

For smaller or biased samples, I partially agree with Hijmans and Graham (2006) that BIOCLIM may underestimate future ranges (agreeing only partially as at the same time, BIOCLIM could continue to overestimate ranges in the ‘B1’ zones). But in these situations, one may also require that more sophisticated algorithms do not extrapolate beyond observed ranges depicted by the grey rectangle in Figure 1 (and see also discussions on SDM transferability as by Charney et al. (2021) for American tree species). Consequently, also the more sophisticated models would then be expected to underestimate the future range. (That more sophisticated models may extrapolate is a phenomenon that we experienced when developing a Climate Change Atlas for Africa (https://atlas.worldagroforestry.org/), where extrapolation from ensemble suitability models tended to result in large area differences if allowed. Realizing this, we included different versions of maps where extrapolations were allowed and where they were not, so that users would be aware of such differences.)

In conclusion, where sample sizes and their representation approximate the true 90% ranges of a particular species, I would expect that the BIOCLIM algorithm can be used as a quick but incomplete filter to identify species that are not suitable (occurring outside the ellipsoid and 90% interval) at a location under baseline or future conditions. The filter is incomplete by correctly identifying unsuitable species but failing to identify a larger set of unsuitable species, with more failures to be expected in situations where correlations between explanatory variables are high.

In the next sections, I will describe the `TreeGOER` database that allows implementation of the BIOCLIM method for nearly 50,000 tree species. I will also illustrate how the database can be used in two global investigations of the effects of climate change.

## 2 COMPILING THE DATABASE

### Presence observations

Presence observations were processed from a recently (within the current decennium, March 2021) compiled data set of 44,267,164 occurrences from the Global Biodiversity Information Facility (GBIF; https://www.gbif.org/) that were collated by Keppel et al. (2021); their appendices S2 and S3) after they had compiled a list of 58,044 tree species that was informed by GlobalTreeSearch version 1.4. These occurrences are available from a GBIF occurrence download (GBIF.org 2021; https://doi.org/10.15468/dl.77gcvq). I selected this data set as GBIF was identified by Keppel et al. (2021) as holding the largest available occurrence data set for tree species.

The data set was subjected to the following series of data quality checks, where records that did not satisfy listed criteria were removed. First, I retained records where information on geographic coordinates (‘decimalLongitude’ and ‘decimalLatitude’) were available. Second, records were removed where information was not available on the ‘basisOfRecord’ and where its value was ‘FOSSIL_SPECIMEN’. Third, records were removed where information on the collection year was not available. Fourth, observations with a collection year prior to 1946 were removed, following the suggestion of the documentation of the `CoordinateCleaner` package (version 2.0-20; Zizka et al., 2019) that GBIF records from and before the second world war period are often very imprecise. Fifth, records were only retained if the geographical coordinates corresponded to a terrestrial 30 arc-seconds (≈ 1 km at the equator) raster cell of the WorldClim 2.1 database (Fick and Hijmans, 2017). Sixth, records were removed if they were flagged by the `CoordinateCleaner::clean_coordinates` function for the ‘capitals’, ‘centroids’, ‘equal’, ‘gbif’, ‘institutions’, ‘zeros’ and ‘duplicates’ tests, thereby flagging records such as those corresponding to country capitals or duplicated coordinate records for the same species.

A final series of data quality checks verified that the country location documented by GBIF in the ‘countryCode’ field corresponded to the country location of Natural Earth (NE) ‘Admin 0 – Countries’ vector layers (first I created an ‘iso3c’ country code variable from the GBIF ‘iso2c’ country code via the `countrycode` package version 1.4.0). I retained all records where the countries matched for the 1:110 million NE layer (https://www.naturalearthdata.com/downloads/110m-cultural-vectors/; version 5.1.1 downloaded in September 2022). From the records where countries did not match, I retained all records where the countries matched for the NE 1:10m layer (https://www.naturalearthdata.com/downloads/10m-cultural-vectors/; version 5.1.1 downloaded in November 2022). By using the small scale layer first, I allowed for some buffers around country boundaries, but such buffers can be considered acceptable given the spatial precision of country GIS layers and locality data (Tack et al., 2022). The NE 1:10 million layer did not show the distribution of certain GBIF country codes as for example ‘GF’ (French Guyana, with corresponding occurrences mainly mapped by NE in the multipolygon with the country code of ‘FRA’, indicating France) or ‘BQ’ (Bonaire, Sint Eustatius and Saba, corresponding in NE to The Netherlands). Recognizing these situations, I also retained occurrence records for the following matches between the GBIF and NE country codes: AX-ALD, BQ-NLD, CX-IOA, EH-SAH, GF-FRA, GP-FRA, MQ-FRA, PS-PSX, RE-FRA, SJ-NOR, SS-SDS and YT-FRA. What was also the case was that some countries and territories mapped by NE had no corresponding country code in the GBIF data set, for example for the separately mapped Somaliland and military bases in Cyprus. I therefore also retained occurrence records for country code matches of: CU-USG, CY-CNM, CY-CYN, CY-WSB and SO-SOL. After realizing that no records had been retained for Namibia, I discovered that the GBIF country code for Namibia of ‘NA’ had been interpreted as missing data when reading in the GBIF data set in `R`. To allow for records from Namibia, I assumed that all records with ‘missing’ GBIF country codes and with occurrences between longitudes 10 and 30 degrees East and latitudes between 15 and 30 South corresponded to observations from Namibia. For these records, the entire process of quality checking was repeated starting from the second step documented above.

### Taxonomy

After data quality checks, I created a master list of the 48,518 unique species names from the retained occurrence records. This result indicated a relatively mild effect of data cleaning by retaining 99.1% of the 48,970 species with at least one observation reported by Keppel et al. (2021) for the same GBIF data set (https://doi.org/10.15468/dl.77gcvq). Via the `WorldFlora` package (version 1.10; Kindt 2020), the master list was standardized to World Flora Online (WFO; Borsch et al., 2020; taxonomic backbone version 2021.12 downloaded from http://www.worldfloraonline.org/downloadData) and, for those species that could not be matched to WFO, to the World Checklist of Vascular Plants (WCVP; Govaerts et al., 2021; taxonomic backbone version 8 downloaded from http://sftp.kew.org/pub/data-repositories/WCVP/). Since these were the same protocols that had been used to create the 2022 version of the Agroforestry Species Switchboard (Kindt et al., 2022), I first directly matched 47,728 species with species that were encountered when preparing the *Switchboard*, followed by using the same matching procedures for the remaining 790 species. The `TreeGOER_Taxonomy` file documents for each of the 48,518 species to which WFO or WCVP taxon they were standardized, further including other details such as the identifier of the taxonomic backbone database and whether the match was made as a direct match or a manual match suggested by fuzzy matching. As some of the submitted names were identified as synonyms, the number of unique standardized names was 48,129, directly corresponding to the species with documented environmental ranges in the `TreeGOER` database.

### Environmental variables

Table S1 in the Supporting Information lists the 51 environmental variables that are covered by `TreeGOER` together with links to references where more details can be obtained about each variable. The bioclimatic, soil and topographic variables can be characterized as abiotic variables and as a consequence, TreeGOER is useful particularly to estimate the ‘abiotically potential range’ of a species (Booth, 2016 and references therein).

A set of 19 bioclimatic layers was obtained at 30 arc-seconds resolution from WorldClim 2.1 (Fick and Hijmans, 2017). Also downloaded from WorldClim 2.1 was the elevation data used when creating the database. I included this elevation data to allow comparisons with elevation ranges documented in other databases or floras, but I strongly advise against using elevation as a predictor variable for suitability in future conditions given its strong correlation with temperature (typically predicted to increase in future climates whereas elevation remains constant; elevation may also be correlated with other factors that affect tree physiology such as frequencies of fog; see also the discussion of ‘indirect variables’ by Guisan and Zimmermann (2000)) – unfortunately this is a mistake that remains to be made in various climate change studies.

With the `envirem` package (version 2.3; Title and Bemmels, 2017), I created an additional set of 18 bioclimatic variables at the same resolution as the WorldClim 2.1 layers (this was also the resolution of the other environmental layers described below). This was done after downloading all historical monthly temperature and rainfall layers from WorldClim 2.1 and creating monthly extraterrestrial solar radiation files for 1985 (at the centre of the 1970 to 2000 period covered by WorldClim 2.1) via `envirem::ETsolradRasters.` The two topographic variables created by the same authors and made available via their ENVIREM website (https://envirem.github.io/) were also included among the environmental layers.

With function `BiodiversityR::ensemble.PET.season` (version 2.14-3; Kindt and Coe, 2006; Kindt, 2018) the maximum climatological water deficit (MCWD) was calculated, corresponding to the dry season with the largest difference between precipitation and potential evapotranspiration (PET). As inputs in the analysis, monthly PET layers were first created via `envirem::monthlyPET`. The calculations involved a similar procedure as used by Chave et al. (2014) who modelled and used this bioclimatic variable to model aboveground biomass of tropical trees, but the `BiodiversityR` procedure allows to distinguish more than one dry season at a location. Zuidema et al. (2022) provided recent evidence for the importance of MCWD in tree growth with data from a pantropical tree-ring network.

Physical and chemical soil properties were obtained via SoilGrids 2.0 (Poggio et al., 2021) by processing 1000 m aggregated data (https://files.isric.org/soilgrids/latest/data_aggregated/1000m/; accessed in November 2022). I obtained eight soil layers that correspond to all chemical and physical soil properties, except for ‘Coarse Fragments (CFVP)’ which I did not include because the Model Efficient Coefficients for this variable were significantly lower (and seeing also the significantly lower R^2^ values reported by Hengl et al., 2017). Using a similar procedure as described by Hannah et al (2020), for each soil variable the average was calculated of strata within the top 1 m (0 - 5 cm, 5 – 15 cm, 15 – 30 cm, 30 – 60 cm and 60 – 100 cm). The average layers were reprojected to the resolution of the bioclimatic layers with `terra::project` (version 1.6-47; Hijmans 2022), using a bioclimatic layer as template.

Also included among the environmental variables were decimal longitude and latitude, directly obtained from the occurrence data set.

### Outlier detection methods

I used two methods for outlier detection based on Tukey’s (1977) fences method and available via `BiodiversityR::ensemble.outliers` (version 2.15-1). In one application of the methodology termed ‘method 1’ here, the default parameters of the function were used for `n_min` (the minimum number of environmental variables required to flag an outlier record) of 5 and `fence.k` (the fence multiplier of the interquartile range) of 2.5. Van Zonneveld et al. (2018) provides the justification for method 1, which was recently also used by van Zonneveld et al. (2023). As a more strict outlier detection method, I also used a ‘method 2’ where `n_min` was set as 2 and `fence.k` as 1.5, the latter as in the original Tukey method. I used both these methods to flag outliers for the 19 bioclimatic variables obtained directly via WorldClim 2.1, not including other bioclimatic layers as these were partially calculated from WorldClim 2.1 data.

### Calculation of global environmental ranges and niche breadths

Global environmental ranges were determined for each species separately. Allowing for different synonyms, all observations for the same standardized species name were pooled together and afterwards duplicated observations from the same 30 arc-seconds grid cell were removed. This procedure reflects a low spatial thinning process whereby some of the spatial sampling biases of species occurrence records can be removed (Aiello-Lammens et al., 2015). Except for longitude and latitude that were obtained directly, values for the other variables were obtained by extracting the 30 arc-seconds raster layers at the locations of the cleaned occurrence data set via `terra::extract`.

After removing outliers flagged by method 1, I calculated the minimum, maximum, median, mean, first quartile and third quartile values via `base::summary`. Quantile estimates at probabilities of 0.05 (Q05) and 0.95 (Q95) were calculated via `Qtools::midquantile` (version 1.5.6; Geraci, 2016). Q05 and Q95 limits are used to define the core distribution of a species in the BIOCLIM method (Lindenmayer et al., 1996; Booth, 2014). Where the number of observations was between 3 and 5000, I calculated the *P* value of a Shapiro-Wilk test of normality via `stats::shapiro.test`.

Relative niche breadth (scaled afterwards as percentage) was calculated for each variable and each species by dividing the species-specific 90% quantile range by the global range; the latter was calculated as the difference between the maximum of all the Q95 values and the minimum of all the Q05 values.

If outlier method 1 identified some outliers, then I also calculated the Q05 and Q95 values for the full data set where outliers were not removed. If outlier method 2 identified outliers, I calculated the Q05 and Q95 values for the data without the respective outliers.

### Statistical software

When developing `TreeGOER` and this article, all data processing was done in `R` (version 4.2.1; R Core Team, 2022). Figures included here were obtained via the `ggplot2` (version 3.3.6; Wickham, 2009) and `sf` (version 1.0-8; Pebesma, 2018) packages. Country outlines in the figures correspond to the Natural Earth 1:110m vector layer used to develop the database.

## 3 DATABASE OVERVIEW AND ACCESS

The `TreeGOER` database documents the bioclimatic ranges for 48,129 species (Table 1). Soil ranges are documented for a smaller number of species (46,608 – 47,635; Table S1 in the Supporting Information), which I attribute to the mask used in SoilGrids that removes built-up, water and glacier areas (Buchhorn et al., 2020). Roughly 45% of species retained 20 observation records or more, 38% retained a minimum of 30 records (this is one of the thresholds identified as a minimum data requirement for species distribution modelling, see introduction), whereas roughly a quarter of the retained species had fewer than 10 records (Table 1). Outlier detection method 2 resulted in lower species counts, with roughly a third of species having 30 records or more and over 3,700 species having no records. Not removing outliers had a relatively mild effect on retained species numbers, with cumulative percentages being maximum one percent higher for bin sizes for 10 observations or more. Given this relatively mild effect, I recommend utilizing `TreeGOER` statistics derived from data sets where outliers were removed. In the remainder of this article, I will therefore use statistics obtained via Method 1. In the below, I will refer to a ‘range’ as defined by the environmental range from Q05 to Q95.

**TABLE 1.** Number of tree species in `TreeGOER` for two different methods of outlier detection (Method 1 and Method 2) and without excluding outliers, calculated for bioclimatic variable bio01. Bin sizes were defined by the number of observations per tree species (N). Values between brackets show cumulative percentages.

| Observations | Method 1 | Method 2 | (All) |
| --- | --- | --- | --- |
| N ≥ 5000 | 256 [0.5] | 234 [0.5] | 258 [0.5] |
| 1000 ≥ N > 5000 | 989 [2.6] | 872 [2.3] | 997 [2.6] |
| 500 ≥ N > 1000 | 1,045 [4.8] | 950 [4.3] | 1,061 [4.8] |
| 200 ≥ N > 500 | 2,748 [10.5] | 2,444 [9.3] | 2,785 [10.6] |
| 100 ≥ N > 200 | 3,613 [18.0] | 3,294 [16.2] | 3,648 [18.2] |
| 50 ≥ N > 100 | 5,084 [28.5] | 4,657 [25.9] | 5,192 [29.0] |
| 30 ≥ N > 50 | 4,704 [38.3] | 4,332 [34.9] | 4,804 [38.9] |
| 20 ≥ N > 30 | 4,009 [46.6] | 3,811 [42.8] | 4,108 [47.5] |
| 10 ≥ N > 20 | 7,065 [61.3] | 6,715 [56.7] | 7,131 [62.3] |
| 5 ≥ N > 10 | 6,253 [74.3] | 6,337 [69.9] | 6,525 [75.9] |
| N = 4 | 2,504 [79.5] | 1,509 [73.0] | 1,976 [80.0] |
| N = 3 | 2,585 [84.9] | 3,577 [80.5] | 2,371 [84.9] |
| N = 2 | 3,150 [91.4] | 2,825 [86.3] | 3,149 [91.4] |
| N = 1 | 4,124 [100.0] | 2,849 [92.3] | 4,124 [100.0] |
| N = 0 | 0 [100.0] | 3,723 [100.0] | 0 [100.0] |

Tables S1 and S2 in the Supporting Information provide information on the distribution of niche breadths for the different variables. For most of the variables, most species and often more than half of them had niche breadths within the 10-25% interval. Notable exceptions for the full data set (Table S1) were bio04, continentality, monthCountByTemp10 (Tmo10), Longitude and Latitude that had most species in the 0-0.5% interval. For species with at least 30 observations (Table S2), only Tmo10 had most species in the 0-0.5% interval, with the second-lowest highest count for latitude in the 2.5-5% bin.

I subdivided the globe in five zones based on the number of months with average temperature above 10 °C (Tmo10) inspired by the Tmo10 thresholds used in the Köppen-Trewartha climate classification system (Belda et al., 2014) to differentiate between tropical, subtropical, temperate, boreal and polar climates whereas I ignored dry climate criteria from that system. Analysing 90% ranges, the vast majority of species (45,274 representing 94.1% of species in `TreeGOER`) occurred in the ‘tropical’ zone were Tmo10 was exactly 12. More than 38,000 species (80.8%) were only observed within this zone (Table 2).

**TABLE 2.** Number of tree species in `TreeGOER` for different global zones defined by the number of months with average temperature > 10 °C (Tmo10), the Climatic Moisture Index (CMI), Latitude (LAT) and Longitude (LON) and where suitability was predicted from 90% ranges of the variable defining the zone. The CMI classification matches dryland zones defined by the aridity index, including dry subhumid (DS), semi-arid (SA), arid (A) and hyperarid (HA) zones. Species were counted separately if their 90% range was contained entirely in the zone (‘endemic’), if they reached their upper distribution limits (Q95) in the zone, if they reached their lower distribution limits (Q05) in the zone or if the limits of the zone was contained entirely with the species 90% range (‘within range’). Values between brackets correspond to species with 30 observations or more. Figure S3 in the Supporting Information provides a zoomable map that shows the global distribution of the different zones.

| Zone | Endemic | Upper limit | Within range | Lower limit |
| --- | --- | --- | --- | --- |
| **Number of months with average temperature > 10 °C** |  |  |  |  |
| Tmo10 = 12 | 38,131 [11,929] | 0 [0] | 7,143 [4,957] | 0 [0] |
| 8 ≤ Tmo10 < 12 | 382 [48] | 1,455 [986] | 2,690 [1,626] | 4,453 [3,331] |
| 4 ≤ Tmo10 < 8 | 354 [140] | 462 [349] | 1,542 [828] | 2,494 [1,779] |
| 1 ≤ Tmo10 < 4 | 13 [0] | 19 [13] | 1,031 [455] | 830 [673] |
| Tmo10 < 1 | 0 [0] | 0 [0] | 668 [261] | 349 [203] |
| **Climatic moisture index** |  |  |  |  |
| CMI ≥ 0.5 | 2,367 [64] | 15,413 [7,431] | 0 [0] | 0 [0] |
| 0 ≤ CMI < 0.5 | 8,899 [829] | 12,458 [6,485] | 5,713 [3,910] | 10,574 [3,982] |
| -0.35 ≤ CMI < 0 | 1,605 [51] | 3,299 [1,707] | 7,190 [4,807] | 10,694 [5,481] |
| -0.5 ≤ CMI < -0.35 (DS) | 201 [1] | 830 [444] | 5,646 [3,876] | 4,578 [2,497] |
| -0.8 ≤ CMI < -0.5 (SA) | 625 [193] | 715 [538] | 581 [416] | 5,697 [3,766] |
| -0.95 ≤ CMI < -0.8 (A) | 198 [47] | 78 [32] | 133 [81] | 1,074 [803] |
| CMI < -0.95 (HA) | 0 [0] | 4 [0] | 4 [1] | 175 [85] |
| **Latitude** |  |  |  |  |
| LAT > 23.5 | 2,880 [1,531] | 2,760 [1,619] | 0 [0] | 0 [0] |
| -23.5 ≤ LAT ≤ 23.5 | 36,943 [11,520] | 3,168 [2,233] | 323 [269] | 2,436 [1,350] |
| LAT < -23.5 | 2,377 [1,536] | 0 [0] | 0 [0] | 3,491 [2,502] |
| **Longitude** |  |  |  |  |
| LON < -30 | 21,581 [9,168] | 0 [0] | 0 [0] | 1,246 [748] |
| -30 ≤ LON ≤ 65 | 8,543 [3,128] | 303 [202] | 943 [546] | 403 [251] |
| LON > 65 | 16,356 [5,144] | 1,345 [797] | 0 [0] | 0 [0] |

A second subdivision of the globe that I implemented was via the Climatic Moisture Index (CMI), where I expanded the dryland classification system based on the aridity index developed by the United Nations Environment Programme (1997) with three humid zones (Table 2). A large number of species (> 17,000) occurred in any of the three humid zones. Nearly 9,000 species exclusively occurred in the zone defined by 0 ≤ CMI < 0.5 where precipitation was equal to double the PET. Fewer species (< 8,000) occurred in the semi-arid zone, mostly species that reached their lowest limits there.

The subdivision based on latitude used the approximate positions of the tropics of Cancer and Capricorn. As for the results with Tmo10, a great majority of species occurred in tropical areas with most of them (nearly 37,000 species) only occurring in that zone (Table 2). Roughly equal amounts of species occurred outside the tropics.

I developed the zones for longitude especially to differentiate between the three main blocks of continental Africa, the Neo-tropics and South-East Asia of tropical moist forests, the most species-rich terrestrial biome of the planet (Couvreur 2015 ; Hagen et al., 2021; Gatti et al., 2022). With the great majority (96.6%) of species occurring exclusively in one of the longitudinal zones (Table 2), a clear pattern emerged of the western zone including the highest number of species and the central zone (including Africa) having substantially lower richness.

Table S3 in the Supporting Information cross tabulates the Tmo10 zones with the other zones. The cross tabulations clearly show that most tree species occur in tropical climates, with one exception for northern latitudes where more species occur in subtropical climates (8 ≤ Tmo10 < 12). Given that `TreeGOER` contains fewer than 49,000 tree species, the crosstabulation of the tropical zone with the longitudinal zones approximated patterns observed for the native distribution of over 58,000 tree species in the *State of the World’s Trees* (BGCI, 2021) reasonably well, where 23,631 species were observed in the Neotropics and 9,237 species in the Afrotropics. A comparison with the BGCI results from the East was more complicated given the latitudinal and longitudinal ranges for Indo-Malaya (13,739 species) and Australasia (7,442 species).

The latitudinal differences clearly follow global latitudinal gradients in biodiversity as for example reviewed by Kinlock et al. (2018) or Nishizawa et al. (2022). Patterns observed in `TreeGOER` also agree with observations that tropical rain forests are the most species-rich terrestrial biomes of the planet and with the “Odd man out” of lower plant and tree diversity in Africa (Couvreur, 2015 ; Raven et al., 2020; Hagen et al., 2021; de Miranda et al., 2022).

The version of the `TreeGOER` database described in this article is publicly archived on Zenodo under a CC-BY 4.0 license so that it can be freely used, shared and modified as long as appropriate credit is given to the database (citing this article and https://doi.org/10.5281/zenodo.7922928). The database is stored as different text files delimited by the pipe “|” character, including the main file with the environmental ranges (TreeGOER_2023.txt), a file with information on taxonomy and standardization methods (TreeGOER_Taxonomy.txt; via the ‘SID’ field more taxonomic details can be easily obtained by linking to the taxonomic backbone databases of WFO or WCVP) and the identifications of the GBIF records involved in calculating the ranges (TreeGOER_GBIFID.txt; to reduce file size, for 126 species with over 10,000 observations, a random subset is provided of 10,000 observations). Also provided are files that show the distribution of species in the Tmo10 and CMI zones with codes reflecting the columns of Table 2; these data can be used for filtering suitable species for different zones.

## 4 CASE STUDIES

To showcase possible applications of the `TreeGOER` database in climate change investigations, I developed two case studies. In both case studies, baseline bioclimatic data were extracted from the raster data used to prepare `TreeGOER`. I used future climate data at 2.5 arc-minutes resolution from the full set of 23 downscaled General Circulation Model (GCM) outputs available for the 2050s and for shared socio-economic pathway 3-7.0 (a higher emissions scenario; Meinshausen et al., 2020) from WorldClim 2.1 (https://www.worldclim.org/data/cmip6/cmip6_clim2.5m.html; accessed in January-March 2023), including ACCESS-CM2, ACCESS-ESM1-5, BCC-CSM2-MR, CanESM5, CanESM5-CanOE, CMCC-ESM2, CNRM-CM6-1, CNRM-CM6-1-HR, CNRM-ESM2-1, EC-Earth3-Veg, EC-Earth3-Veg-LR, GFDL-ESM4, GISS-E2-1-G, GISS-E2-1-H, INM-CM4-8, INM-CM5-0, IPSL-CM6A-LR, MIROC-ES2L, MIROC6, MPI-ESM1-2-HR, MPI-ESM1-2-LR, MRI-ESM2-0 and UKESM1-0-LL. Estimates for bioclimatic variables directly available from WorldClim were calculated as median values after extracting the 23 separate layers at the point locations of the case studies. The extended set of `envirem` variables was calculated at point locations for the different GCMs via `BiodiversityR::ensemble.envirem.masterstack`, `BiodiversityR::ensemble.envirem.solradstack`, and `BiodiversityR::envirem`.run. Prior to these calculations, monthly layers of extraterrestrial solar radiation were generated for 2050 via `envirem::ETsolradRasters`. Similar to the bioclimatic data from WorldClim, I calculated medians afterwards across the 23 GCMs.

I selected a subset of environmental variables for the case studies that included bio01 (mean annual temperature), bio12 (annual precipitation), bio05 (maximum temperature of the warmest month), bio06 (minimum temperature of the coldest month), bio16 (precipitation of the wettest quarter), bio17 (precipitation of the driest quarter), climaticMoistureIndex (CMI), monthCountByTemp10 (Tmo10) and growingDegDays5 (GDD5). I selected these variables as they corresponded to bioclimatic variables used previously for BIOCLIM modelling of the future climatic adaptability of tree species by Nogués-Bravo et al. (2014) and Booth (2016) (GDD5, bio06 and a moisture index) or otherwise used by Esperon-Rodriguez et al. (2022) (bio01, bio05, bio06, bio12 and bio17). I expanded the set with bio16 to add a humid period variable to the dry period variable of bio17 analogous to having the cold period and warm period variables of bio05 and bio06. I also added Tmo10 as it was used for zonation when describing `TreeGOER` above, but this variable was redundant for case study 2.

I used a subset of the `TreeGOER` database where 5% and 95% statistics have been calculated from 20 or more observations; these were available for 22,448 species (Table 1). Esperon-Rodriguez et al. (2022) used the same criterion to create a species subset for their study. From my discussion of Figure 1 in the introduction, my assumption would be that the results obtained for the case studies are conservative, but this assumption also implies that the available records were representative for the environmental ranges of most species for the selected variables, and that these variables characterize the suitable niche well (see a discussion by Santini et al., 2021).

### Case study 1: Testing the effect of climate change for urban locations

Esperon-Rodriguez et al. (2022) conducted a global study on the climate change vulnerability of 3,129 urban trees and shrubs. These authors also used median values for the 2050s, albeit for a different high emissions scenario (RCP 6.0) and a set of ten GCM projections specific for urban climates (Zhao et al., 2021). Their analyses were based on 5% and 95% quantiles as well, but they selected 95% thresholds for temperature-based bioclimatic variables and 5% thresholds for precipitation-based ones (see above which variables they selected). A fundamental part of their analyses is that their estimates were done for each city separately for documented city-species observations (Ossola et al., 2020). Where the results that they present are for 164 cities, I excluded three cities as the geographical coordinates were outside the baseline 30 arc-seconds raster terrestrial coverage. Given these differences in methodologies, my main objective here, besides showcasing that straightforward climate change analyses via `TreeGOER` can be conducted, therefore was to check whether still similar results would be obtained. I especially checked the Esperon-Rodriguez et al. (2022) finding that cities at low latitudes would be more vulnerable to climate change.

I estimated future species richness by two different methods. The first method (‘filter 1’) identified all species suitable to the future climate at the city location. The second method (‘filter 2’) first filtered species suitable to the baseline climate and then proceeded to further filter those species that remained suitable to the future climate. The difference between the two methods can be understood as allowing species to expand their ranges to occupy previously unoccupied sites, or not allowing migration to track suitable climatic conditions (Boisvert-Marsh and de Blois, 2021; Lima et al., 2022).

With filter 1, locations furthest from the equator often were predicted to have larger proportions of species richness in the future climate (FP; Figure 2b; Data S5 in the Supporting Information). With filter 2, every city was predicted to lose some species, but generally more so at lower latitudes (Figure 2c).

**FIGURE 2.**
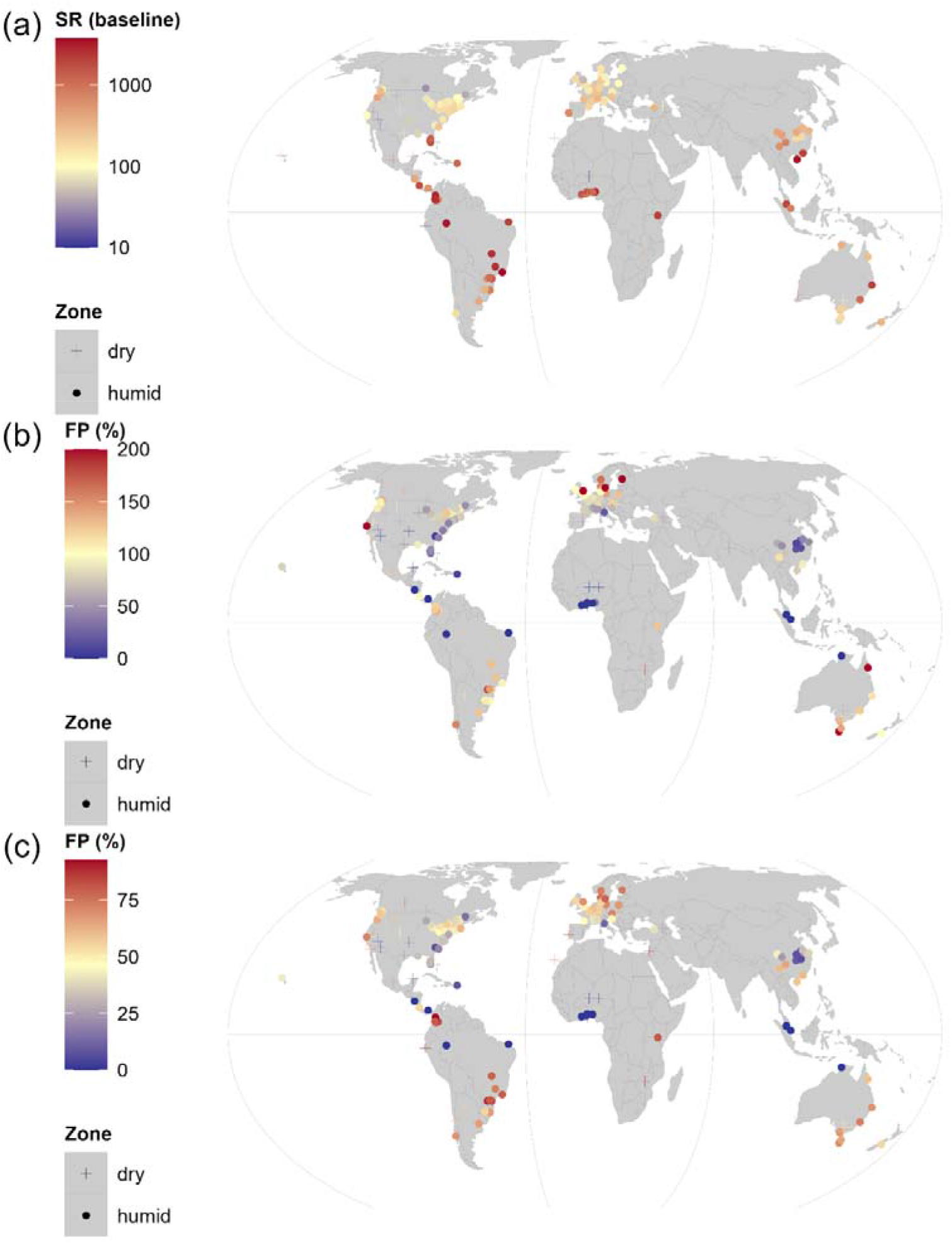
Predicted species richness (SR) of suitable tree species in the baseline climate (a) and (b-c) predicted future proportions (FP) suitable in the future climate for 161 locations of cities (Data S5 in the Supporting Information). Panel (b) shows results where all suitable species were included, with proportions above 200% depicted as 200% for six locations. Panel (c) shows results where only species suitable in the baseline climate at the city location were investigated for future suitability, a “no migration” scenario. Dryland locations were defined by a CMI < -0.35 conform global dryland definitions. Map lines obtained from a Natural Earth 1:110m vector layer (see methods) delineate study areas and do not necessarily depict accepted national boundaries. The maps were created in `R` with Equal Earth projection.

**FIGURE 3.**
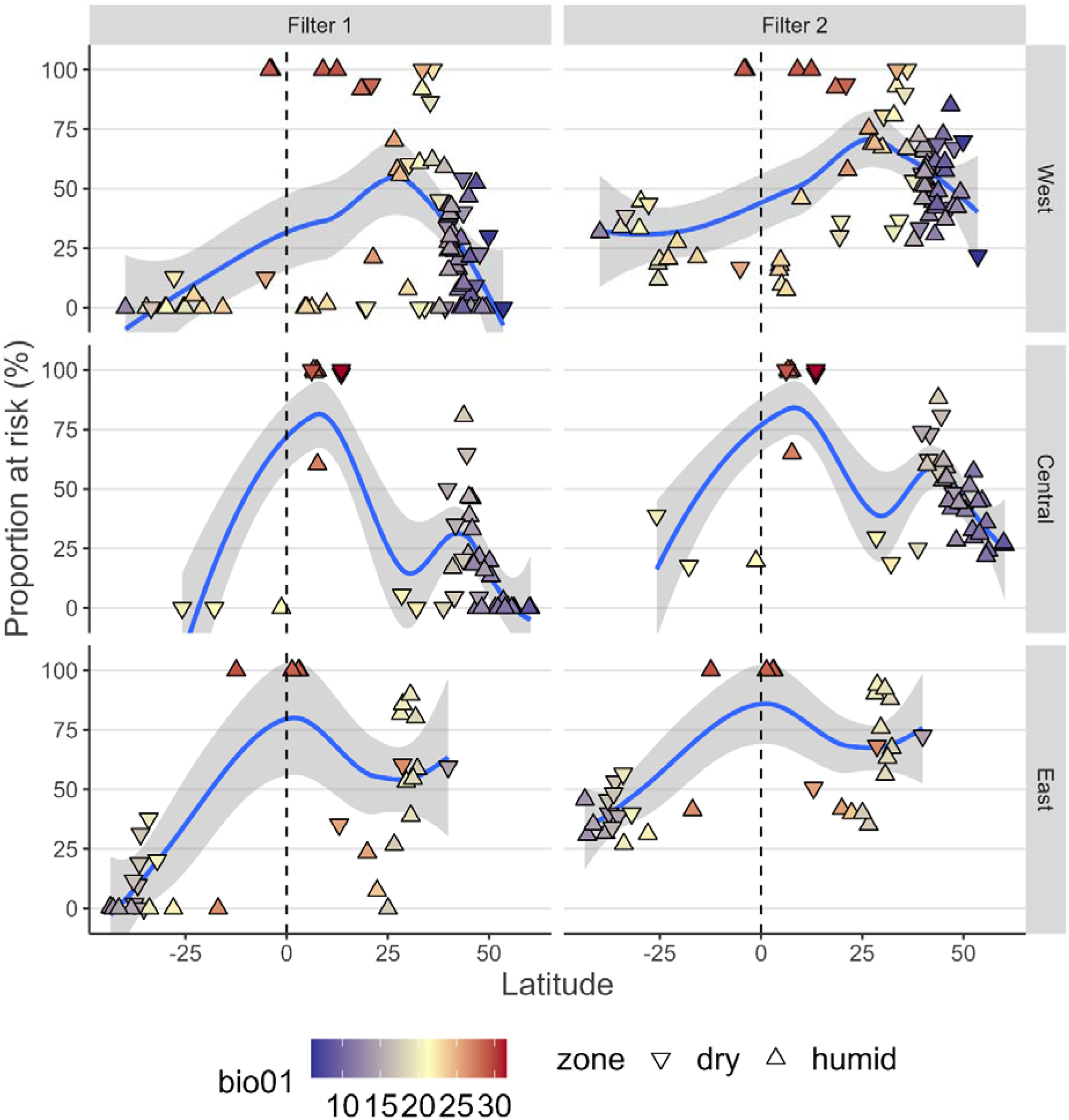
Predicted proportions of species at risk for 161 locations of cities (Figure 4; the Central zone has longitudes between -30 and 65). With filter 1, all suitable species were included. With filter 2, only species suitable in the baseline climate at the city location were investigated. The colour scale corresponds to bio01, the mean annual temperature. Dryland locations were defined by a CMI < -0.35 conform global dryland definitions. Smoothed regression curves were added via `ggplot2::geom_smooth` (version 3.3.6) with the `loess` method. Dashed vertical reference lines corresponds to the equator.

The results shown here (Figure 3) confirm the general pattern of increased vulnerability at lower latitudes that Esperon-Rodriguez et al. (2022) had documented. When investigating vulnerability (calculated as `100 – min(c(FP, 100))`) separately for three longitudinal zones, however, in the west the highest vulnerability manifested at mid-latitudes in the North. What I also observed was that cities near the equator with lower annual mean temperatures (a pattern that can be explained by their higher altitudes; see also Liang et al. (2022) and their discussion on the annual mean temperature as dominant predictor of global tree species richness patterns), such as Nairobi for the central zone and Bogota in the west, had a significantly lower risk. An effect of the moisture index was not immediately obvious with dryland and humid locations showing the full range of risks. What should not be forgotten both for the results of Esperon-Rodriguez et al. (2022) and those shown here is that the latitudinal pattern only explains a relatively small part of the variation among cities.

### Case study 2: Testing the effect of climate change for tropical locations

In the second case study, I focused the climate change modelling on tropical areas (Tmo10 = 12) where I also excluded arid and hyperarid zones (CMI < -0.8; Table 2). This is the global area that is richest overall in plant and tree species, especially in humid areas (Couvreur 2015 ; Keppel et al. 2021; Gatti et al., 2022; Table S4 in the Supporting Information). I randomly sampled 2000 locations in this study area via `dismo::randomPoints` (version 1.3-9; Hijmans et al., 2022; the function uses a weighted sampling adjustment for longitude-latitude coordinate systems). Besides mapping overall patterns and differences of tree species richness in the baseline and future climate, I focused on patterns related to the moisture index as one of the explanatory variables for differences in species richness (this bioclimatic variable explains differences in total species richness between zones as documented and discussed in the previous section (Tables 2 and S4), but no obvious pattern had emerged in the first case study). The moisture index can also be linked to inter- and intra-specific variations in carbon sequestration potential (Jucker et al., 2022).

Results for the baseline and future climate are available from Table S6 in the Supplementary Information and are shown in figures 4 and 5. Under changed climate conditions for the middle of the 21^st^ century and the higher emissions scenario, significant effects of climate change can be observed and especially so in South America (Figure 4 bc). These patterns can be seen clearly when comparing results for the different longitudinal zones separately (Figure 5). Strong declines in species richness can be seen in the west and especially so in humid zones with CMI > 0 (Figure 5). Also in the east, an overall trend of strong decline in species richness can be observed. The trend is less pronounced in central Africa, which can be partially explained by a larger fraction of species being suitable in this zone in future conditions.

**FIGURE 4.**
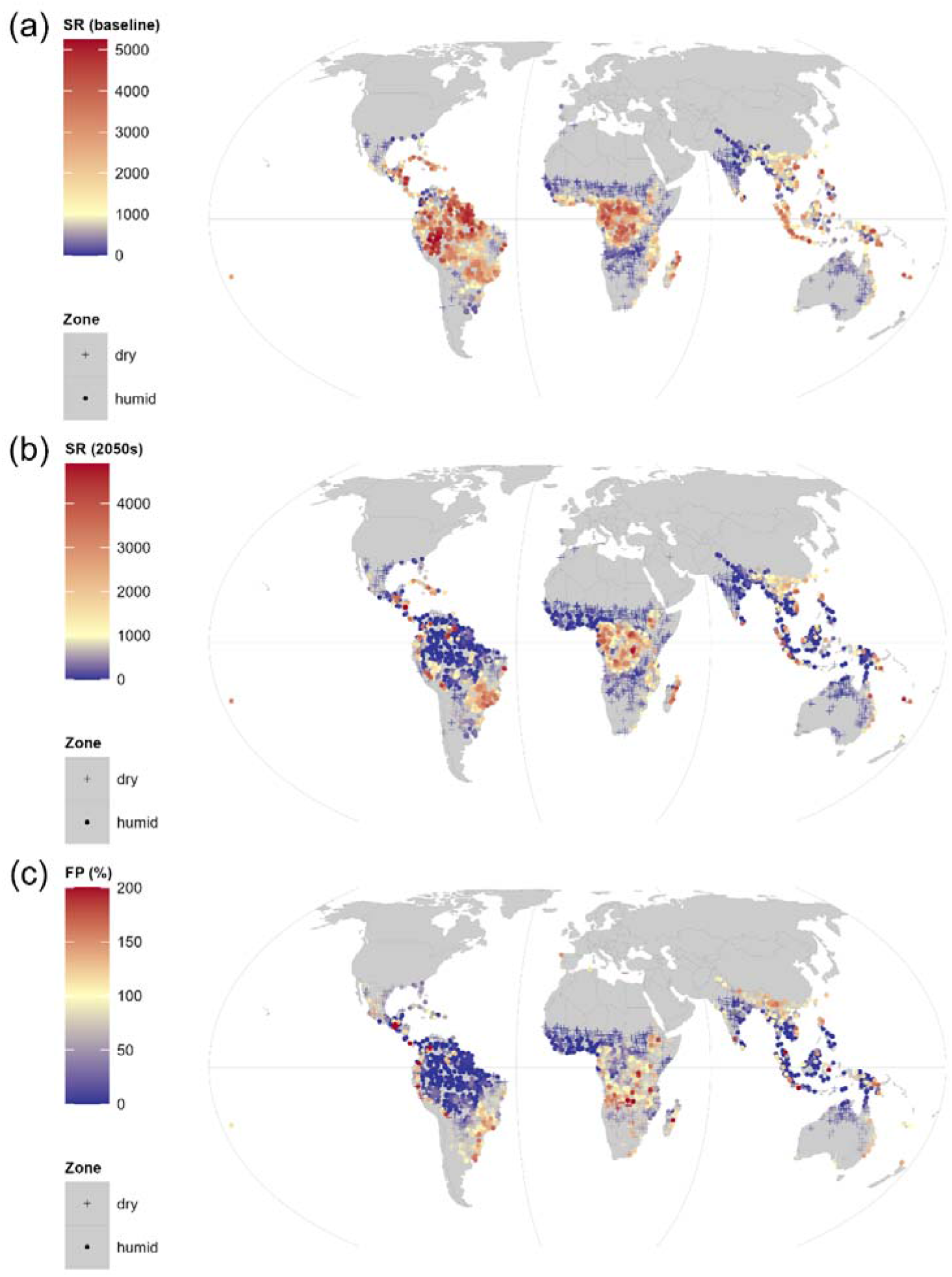
Predicted species richness (SR) of suitable tree species in the (a) baseline and (b) future climate for 2000 randomly selected locations in tropical areas that exclude (hyper-)arid zones (Tmo10 = 12 and CMI > -0.8; Data S6 in the Supporting Information). Panel (c) shows the future SR as a proportion of the baseline SR, with 20 locations with proportions above 200% depicted as 200%. Dryland locations were defined by a CMI < -0.35 conform global dryland definitions. Map lines obtained from a Natural Earth 1:110m vector layer (see methods) delineate study areas and do not necessarily depict accepted national boundaries. The maps were created in `R` with Equal Earth projection.

**FIGURE 5.**
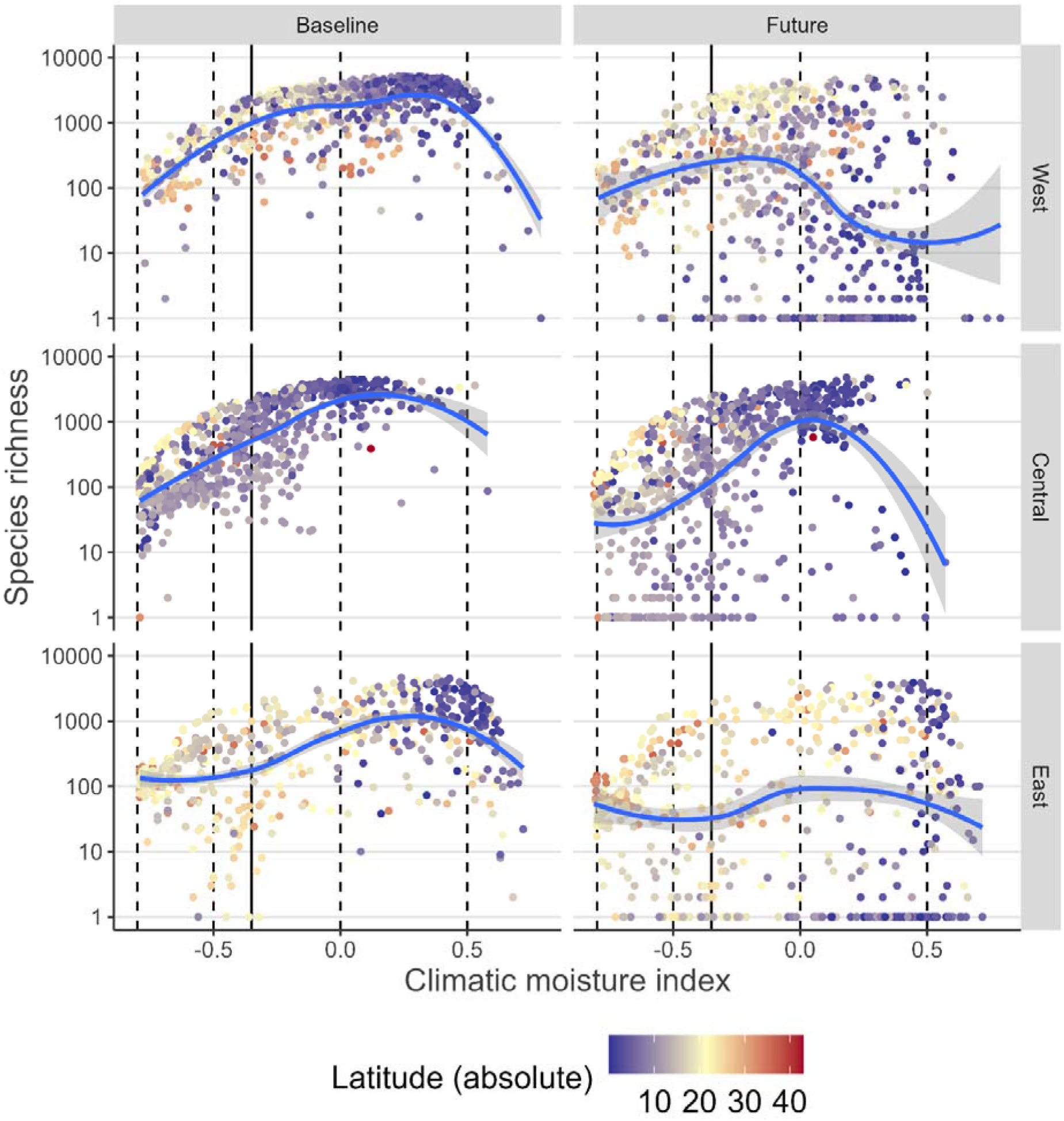
Predicted species richness of suitable tree species in the baseline and future climate for three longitudinal zones for 2000 randomly selected locations in tropical areas that exclude (hyper-)arid zones (see Figure 2 and Figure S3 in the Supporting Information; the Central zone has longitudes between -30 and 65). Shown richness values were transformed by adding 1, hence values of 1 correspond to a predicted species richness of 0. Smoothed regression curves were added via `ggplot2::geom_smooth` (version 3.3.6) with the `loess` method. Full vertical reference lines correspond to the CMI = -0.35 upper boundary of dryland zones, with dashed lines delimiting other CMI zones from Table 3.

Figure 5a clearly suggests an effect of water availability, with dryland locations typically having lower predicted species richness. Figure 5c suggests that various dryland locations may become suitable to a larger range of species, for example in dryland areas in southern and eastern Africa, but not in west Africa.

The highest species richness observed in the baseline climate was 5,252 in a Brazilian location in the western Amazon (longitude: -72.4042; latitude: -7.9042). All ten observations of richness above 5,000 were in Brazil. The highest observation of 4,635 species outside South America was in the Philippines at rank 42. The highest species richness was in a similar range between 5,000 and 6,000 species as shown by Keppel et al. (2021) across 463 geographic regions. What is interesting is that the results shown here are based on point estimates (α-diversity), whereas the Keppel et al. (2021) results refer to regional statistics (γ- diversity). Cai et al. (2023) who recently conducted a sophisticated modelling exercise to predict plant diversity arrived at maximum estimates near 5,000 (and also important to note is that the study identified bioclimatic variables as the most important drivers of diversity; see also Keil and Chase (2019) and Liang et al. (2022) with similar observations for global studies of tree diversity). As my calculations had involved all candidate species regardless their native distribution and I had expected that this could partially explain differences with the other studies, I conducted an analysis where only species native to a particular country were investigated for baseline and future suitability (Figures S6 and S7 in the Supplementary Information; data on native country distribution was obtained from GlobalTreeSearch; Beech et al., 2017). Generally, the same patterns can be seen for native species as with the full set of species, but with maximum species richness values now below 2,500.

## 5 FUTURE DEVELOPMENTS

Looking ahead, when sufficient new observations become available in GBIF and particularly for species that are not yet covered, had low numbers of observations or with remaining large biases in geographical coverage (Meyer et al., 2016), `TreeGOER` could be expanded with this new information. With new plants that continue to be described (Raven et al., 2020) and estimates that the total number of tree species may be larger than 73,000 (Cazzolla Gatti et al., 2022), it is likely that coverage of `TreeGOER` could be expanded substantially. Potentially new observations would become available from locations of novel climate conditions (Williams and Jackson; 2007), what would benefit estimations of future geographical ranges. When updating `TreeGOER` for the new records, taxonomical changes should also be considered.

Another reason to update `TreeGOER` would be when new versions of WorldClim or SoilGrids were released, as differences between previous versions are known to result in differences in climate change predictions (Cerasoli et al., 2022). One of the reasons that WorldClim might need a future revision is that it contains some anomalies for bioclimatic variables that combine temperature and precipitation data (Booth, 2022).

It is likely that I may expand `TreeGOER` with bioclimatic ranges inferred from CHELSA (Climatologies at high resolution for the earth’s land surface areas) as this data set contains several unique bioclimatic variables and also used alternative methods of statistical downscaling compared to the ones used by WorldClim, such as estimating orographic and wind effects on precipitation (Karger et al., 2017; https://chelsa-climate.org/bioclim/). What could also be an important addition are observation data from a proposed global tree trial database (Booth 2023), especially if it was not straightforward to extract these data directly from such database. As fine-scale gradients in soil conditions have been shown to affect species distributions (e.g. Bartholomew et al., 2022), it may be wise to compare quantiles of soil variables observed in `TreeGOER` with those obtained from high resolution soil surveys – provided databases are available that document the full range of suitable soil conditions inferred by detailed sampling.

## AUTHORS’ CONTRIBUTIONS

R.K. conceived the idea and compiled the `TreeGOER` database by the processes documented here. He also created the different drafts of the manuscript, including all its analyses and figures.

## ACKNOWLEDGEMENTS

I am indebted to the countless individuals and organizations that were involved in collecting and availing species occurrence data available via the GBIF occurrence download (GBIF.org 2021; https://doi.org/10.15468/dl.77gcvq). I greatly appreciate the technical feedback provided by Ian Dawson, Lars Graudal, David Bartholomew, Jens-Peter Lillesø, Fabio Pedercini and Rhett Harrison which helped me improve the paper, the writing assistance provided by Ian Dawson and for the acquisition of funds and other support provided by Ramni Jamnadass and Paul Smith. R.K. was supported by the Darwin Initiative to project DAREX001 of *Developing a Global Biodiversity Standard certification for tree-planting and restoration*, by Norway’s International Climate and Forest Initiative through the Royal Norwegian Embassy in Ethiopia to the *Provision of Adequate Tree Seed Portfolio* project in Ethiopia and by the Green Climate Fund through the IUCN-led *Transforming the Eastern Province of Rwanda through Adaptation* project.

## DATA AVAILABILITY STATEMENT

The version of the `TreeGOER` database described in this paper is permanently archived on Zenodo (https://doi.org/10.5281/zenodo.7922928). This repository contains a metadata file describing each field of the different files of the database. `R` code and ancillary data needed to replicate the case studies presented in this paper is available in the Supporting Information.

## Notes

https://doi.org/10.5281/zenodo.7922927

### Competing Interest Statement

The authors have declared no competing interest.

