## Supporting Information 2 for "TreeGOER: a database with globally observed environmental ranges for 48,129 tree species"

**FIGURE S3.** Highly-zoomable maps showing global zones defined by the number of months with average temperature > 10 °C (Tmo10; top) and the Climatic Moisture Index (CMI; bottom). The CMI classification incorporates the dryland zones of dry subhumid (-0.5 ≤ CMI < -0.35), semi-arid (-0.8 ≤ CMI < -0.5), arid (-0.95 ≤ CMI < -0.8) and hyperarid (CMI < -0.95) zones. Latitudinal and longitudinal reference lines correspond to the additional global zonation systems used for Table 2 and Table S4 in the manuscript. These maps were created from 30 arc-seconds baseline (1970-2000) monthly precipitation and minimum, maximum and average temperature raster layers obtained from WorldClim 2.1 and processed by the `envirem` package. Map lines obtained from a Natural Earth 1:110m vector layer (see methods) delineate study areas and do not necessarily depict accepted national boundaries. The map was created in R via the `terra` package.

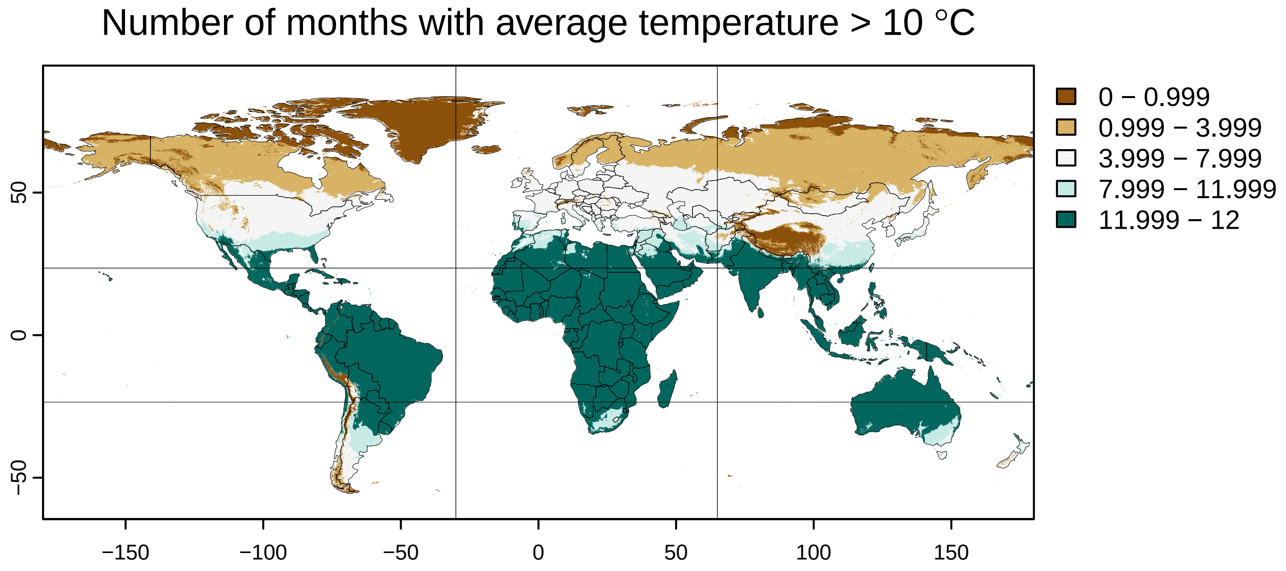

Number of months with average temperature > 10 °C
0 − 0.999
0.999 − 3.999
3.999 − 7.999
7.999 − 11.999
11.999 − 12
50
0
−50
−150
−100
−50
0
50
100
150


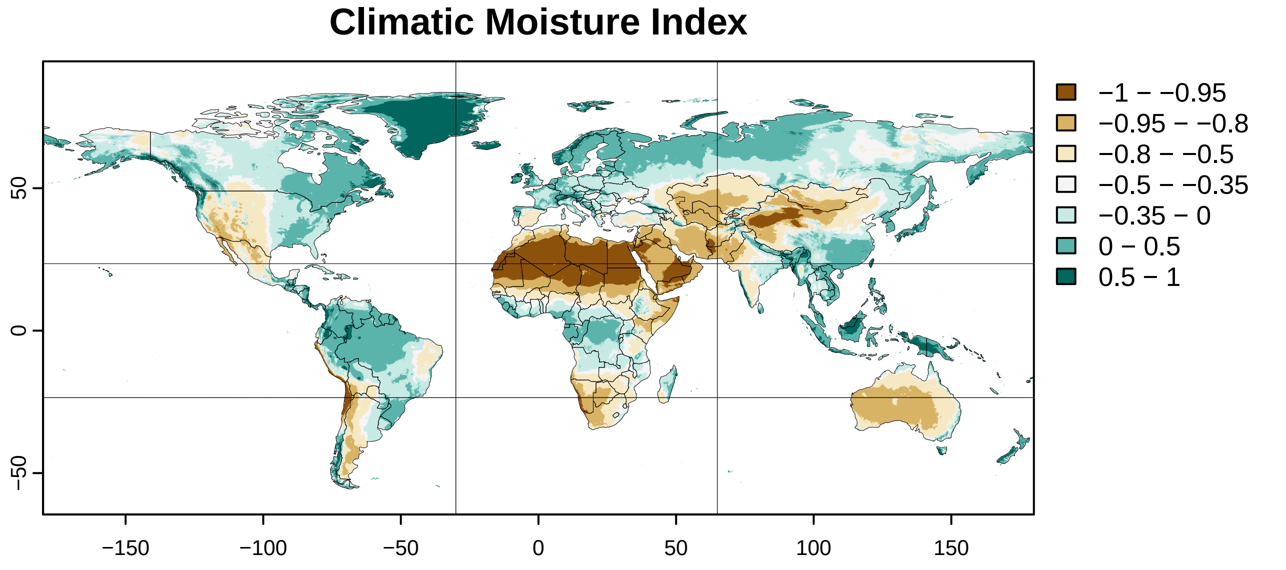

Climatic Moisture Index
−1 − −0.95
−0.95 − −0.8
−0.8 − −0.5
−0.5 − −0.35
−0.35 − 0
0 − 0.5
0.5 − 1
50
0
−50
−150
−100
−50
0
50
100
150

**FIGURE S7**

Predicted species richness (SR) of suitable native tree species in the (a) baseline and (b) future climate for 2000 randomly selected locations in tropical areas that exclude (hyper-)arid zones (Tmo10 = 12 and CMI > -0.8; Data S6 in the Supporting Information). Panel (c) shows the future SR as a proportion of the baseline SR, with 36 locations with proportions above 200% depicted as 200%. Dryland locations were defined by a CMI < -0.35 conform global dryland definitions. Map lines obtained from a Natural Earth 1:110m vector layer (see methods) delineate study areas and do not necessarily depict accepted national boundaries. The maps were created in R with Equal Earth projection.

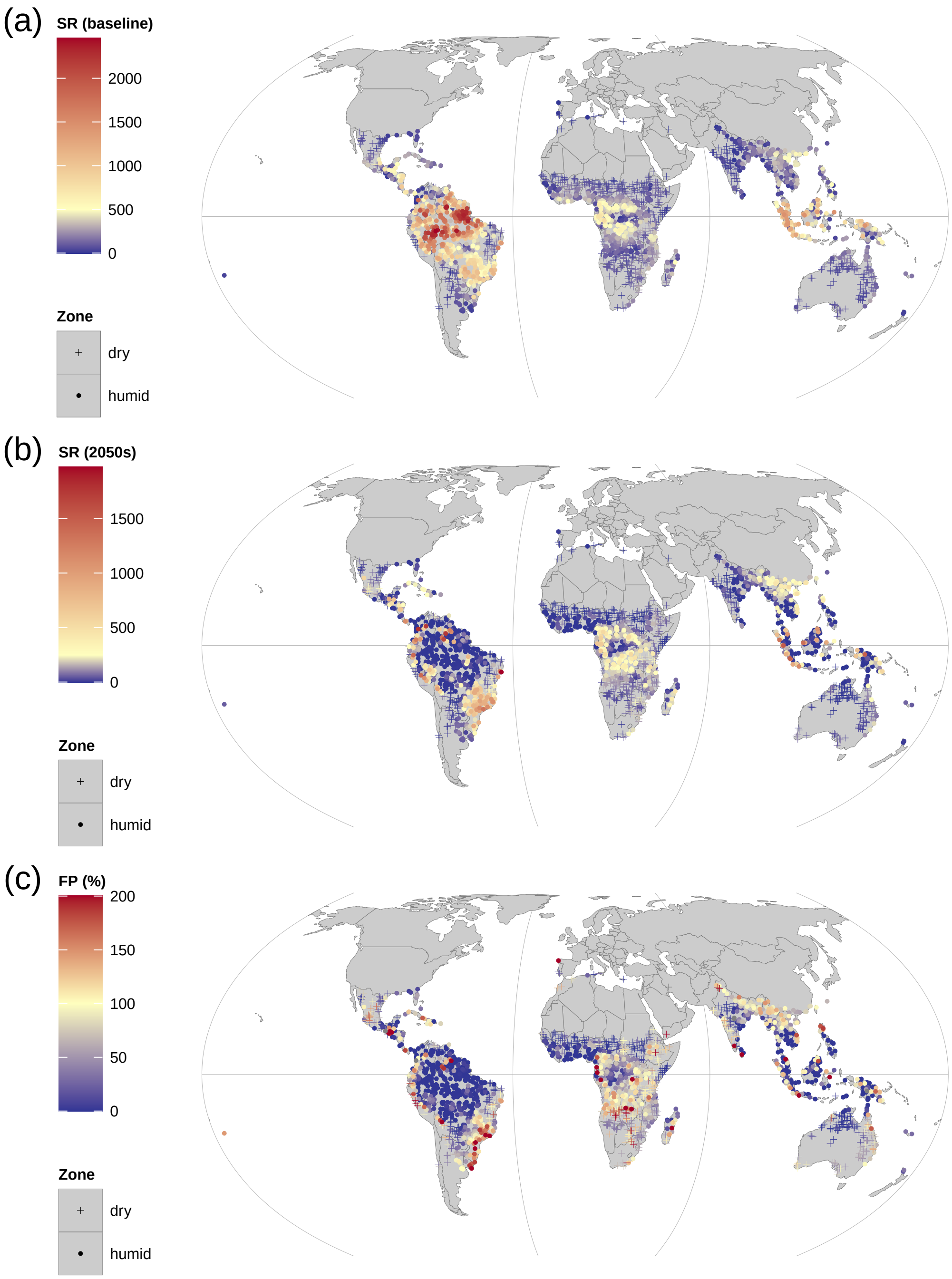

(a)
SR (baseline)
2000
1500
1000
500
0
Zone
dry
humid
(b)
SR (2050s)
1500
1000
500
0
Zone
dry
humid
(c)
FP (%)
200
150
100
50
0
Zone
dry
humid

**FIGURE S8**

Predicted species richness of suitable native tree species in the baseline and future climate for three longitudinal zones for 2000 randomly selected locations in tropical areas that exclude (hyper-)arid zones (see Figure S7 in the Supporting Information; the Central zone has longitudes between -30 and 65). Shown richness values were transformed by adding 1, hence values of 1 correspond to a predicted species richness of 0. Smoothed regression curves were added via `ggplot2::geom_smooth` (version 3.3.6) with the `loess` method. Full vertical reference lines correspond to the CMI = -0.35 upper boundary of dryland zones, with dashed lines delimiting other CMI zones from Table 3.

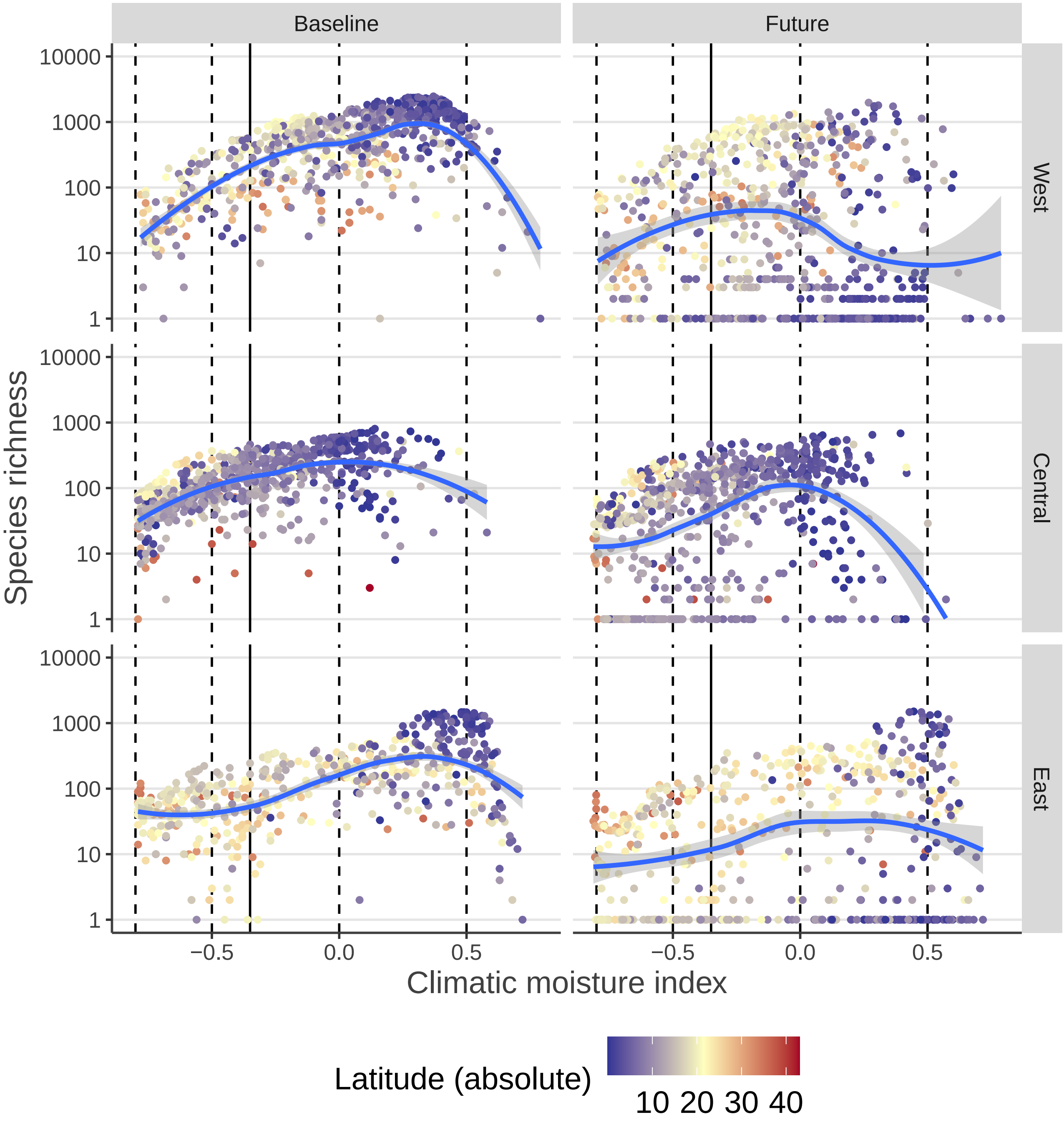

Baseline
Future
West
Central
East
Species richness
10000
1000
100
10
1
Climatic moisture index
−0.5
0.0
0.5
Latitude (absolute)
10 20 30 40
