## Supporting Information 3 for "TreeGOER: a database with globally observed environmental ranges for 48,129 tree species"

### Supplementary script

Roeland KINDT

2023-05-08

**7 Session information** **15**

### 1 Introduction

The scripts shown here are the core scripts for calculating the number of suitable tree species for the two case studies.

- For case study 1, a global data set with city locations was used.
- For case study 2, a random subset was made in tropical locations.

### 2 Libraries needed

```
library(data.table)
library(readxl)
```

### 3 Globally observed environmental ranges in TreeGOER

#### 3.1 Load the data

Load the main file of the TreeGOER database after downloading it to a local folder.

```
# TreeGOER.file <- choose.files()
TreeGOER.file <- "C://Data/TreeGOER_2023.txt"
ranges <- data.frame(fread(file=TreeGOER.file, sep="|", encoding="UTF-8"))
nrow(ranges)
```

```
## [1] 2450209
```

```
head(ranges)
```

```
##            species   var n    MIN   QRT1 MEDIAN   MEAN   QRT3    MAX    Q05
## 1 Abarema abbottii bio01 3  22.84  24.15  25.45  24.67  25.59  25.72  22.84
## 2 Abarema abbottii bio02 3   9.28   9.39   9.50   9.52   9.64   9.78   9.28
## 3 Abarema abbottii bio03 3  73.01  74.20  75.40  74.60  75.40  75.41  73.01
## 4 Abarema abbottii bio04 3 117.66 118.86 120.06 124.97 128.62 137.19 117.66
## 5 Abarema abbottii bio05 3  29.40  30.45  31.50  30.87  31.60  31.70  29.40
## 6 Abarema abbottii bio06 3  16.00  17.45  18.90  18.10  19.15  19.40  16.00
##      Q95 breadthPercentage SHAP outliers n.out Q05.out Q95.out outliers.class
## 1  25.72             6.348 0.16    FALSE    NA      NA      NA          FALSE
## 2   9.78             3.014 0.87    FALSE    NA      NA      NA          FALSE
## 3  75.41             2.957 0.01    FALSE    NA      NA      NA          FALSE
## 4 137.19             0.872 0.22    FALSE    NA      NA      NA          FALSE
## 5  31.70             5.671 0.15    FALSE    NA      NA      NA          FALSE
## 6  19.40             4.877 0.26    FALSE    NA      NA      NA          FALSE
##   n.class Q05.class Q95.class
## 1      NA        NA        NA
## 2      NA        NA        NA
## 3      NA        NA        NA
## 4      NA        NA        NA
## 5      NA        NA        NA
## 6      NA        NA        NA
```

```
summary(ranges)
```

```
##    species              var                  n                 MIN
##  Length:2450209     Length:2450209     Min.   :     1.0   Min.   :-2219.67
##  Class :character   Class :character   1st Qu.:     4.0   1st Qu.:   12.08
##  Mode  :character   Mode  :character   Median :    17.0   Median :   45.30
##                                        Mean   :   244.7   Mean   :  395.80
##                                        3rd Qu.:    61.0   3rd Qu.:  188.10
##                                        Max.   :211181.0   Max.   :10867.40
##
##       QRT1              MEDIAN             MEAN              QRT3
##  Min.   :-1999.20   Min.   :-1969.8   Min.   :-1980.75   Min.   :-1951.3
##  1st Qu.:   17.99   1st Qu.:   20.0   1st Qu.:   20.32   1st Qu.:   21.7
##  Median :   61.25   Median :   71.0   Median :   73.72   Median :   81.3
##  Mean   :  476.92   Mean   :  517.4   Mean   :  519.34   Mean   :  559.7
##  3rd Qu.:  267.70   3rd Qu.:  302.6   3rd Qu.:  311.21   3rd Qu.:  341.0
##  Max.   :10867.40   Max.   :10903.5   Max.   :10867.40   Max.   :10924.9
##
##       MAX                Q05                Q95          breadthPercentage
##  Min.   :-1939.71   Min.   :-2043.72   Min.   :-1939.71   Min.   :  0.000
##  1st Qu.:   24.03   1st Qu.:   14.44   1st Qu.:   23.32   1st Qu.:  3.053
##  Median :  102.00   Median :   50.82   Median :   93.55   Median :  9.283
##  Mean   :  661.41   Mean   :  423.07   Mean   :  622.35   Mean   : 12.308
##  3rd Qu.:  443.90   3rd Qu.:  215.08   3rd Qu.:  405.22   3rd Qu.: 17.835
##  Max.   :11426.50   Max.   :10867.40   Max.   :11122.12   Max.   :100.000
##
##       SHAP            outliers           n.out            Q05.out
##  Min.   :-1.00000   Mode :logical   Min.   :     1.0   Min.   :-2041.0
##  1st Qu.: 0.00000   FALSE:1535244   1st Qu.:    16.0   1st Qu.:   12.5
##  Median : 0.01000   TRUE :914965    Median :    45.0   Median :   44.5
##  Mean   :-0.01191                   Mean   :   588.1   Mean   :  384.5
##  3rd Qu.: 0.23000                   3rd Qu.:   154.0   3rd Qu.:  180.8
##  Max.   : 1.00000                   Max.   :216647.0   Max.   :10100.9
##                                     NA's   :1535244    NA's   :1535244
##     Q95.out       outliers.class     n.class          Q05.class
##  Min.   :-1722.3   Mode :logical   Min.   :     0.0   Min.   :-2051.25
##  1st Qu.:   24.3   FALSE:215828    1st Qu.:     5.0   1st Qu.:   15.60
##  Median :  103.0   TRUE :2234381   Median :    17.0   Median :   53.41
##  Mean   :  649.6                   Mean   :   234.6   Mean   :  441.18
##  3rd Qu.:  432.9                   3rd Qu.:    60.0   3rd Qu.:  225.90
##  Max.   :11023.6                   Max.   :181961.0   Max.   :10864.00
##  NA's   :1535244                   NA's   :215828     NA's   :216132
##    Q95.class
##  Min.   :-1877.70
##  1st Qu.:   23.28
##  Median :   93.00
##  Mean   :  612.10
##  3rd Qu.:  394.90
##  Max.   :11098.10
##  NA's   :216132
```

```r
vars.unique <- unique(ranges$var)# environmental variables covered by TreeGOER
data.frame(Count=c(1:length(vars.unique)),
           Variable=vars.unique)
```

```
##    Count                   Variable
## 1      1                      bio01
## 2      2                      bio02
## 3      3                      bio03
## 4      4                      bio04
## 5      5                      bio05
## 6      6                      bio06
## 7      7                      bio07
## 8      8                      bio08
## 9      9                      bio09
## 10    10                      bio10
## 11    11                      bio11
## 12    12                      bio12
## 13    13                      bio13
## 14    14                      bio14
## 15    15                      bio15
## 16    16                      bio16
## 17    17                      bio17
## 18    18                      bio18
## 19    19                      bio19
## 20    20                  annualPET
## 21    21 aridityIndexThornthwaite
## 22    22    climaticMoistureIndex
## 23    23           continentality
## 24    24                embergerQ
## 25    25           growingDegDays0
## 26    26           growingDegDays5
## 27    27            maxTempColdest
## 28    28           meanTempColdest
## 29    29           meanTempWarmest
## 30    30            minTempWarmest
## 31    31        monthCountByTemp10
## 32    32         PETColdestQuarter
## 33    33          PETDriestQuarter
## 34    34            PETseasonality
## 35    35         PETWarmestQuarter
## 36    36         PETWettestQuarter
## 37    37           thermicityIndex
## 38    38                      MCWD
## 39    39                      bdod
## 40    40                       cec
## 41    41                      clay
## 42    42                  nitrogen
## 43    43                     phh2o
## 44    44                      sand
## 45    45                      silt
## 46    46                       soc
## 47    47                   topoWet
## 48    48                       tri
```

```
## 49    49                    elev
## 50    50                     LON
## 51    51                     LAT
```

For the case studies, we limit the species to those where a minimum of 20 observations were retained. We use the data that where outliers were removed with the default methodology (method 1 as described in the manuscript).

```
ranges <- ranges[ranges$n > 19, ]
nrow(ranges)
```

```
## [1] 1138971
```

```
length(unique(ranges$species))
```

```
## [1] 22448
```

#### 3.2 Ranges for the selected bioclimatic variables

A new data.frame needs to be created with different columns for the ranges (5% and 95%) of the chosen bioclimatic variables.

```
focal.vars <- c("bio01", "bio12",
                "climaticMoistureIndex", "monthCountByTemp10", "growingDegDays5",
                "bio05", "bio06", "bio16", "bio17")
```

```
focal.var <- focal.vars[1]
focal.ranges <- ranges[ranges$var == focal.var, c("species", "n", "Q05", "Q95")]
names(focal.ranges)[3:4] <- paste0(focal.var, "_", names(focal.ranges)[3:4])
ranges.lookup <- focal.ranges
```

When adding columns for the second, third, … bioclimatic variables, a check is done that these data represent the same species. For some variables such as soil variables, it may be necessary to modify the process (e.g. using the dplyr::left_join function) to ensure that species names match.

```
for (i in 2:length(focal.vars)) {
  focal.var <- focal.vars[i]
  focal.ranges <- ranges[ranges$var == focal.var, c("species", "n", "Q05", "Q95")]
  names(focal.ranges)[3:4] <- paste0(focal.var, "_", names(focal.ranges)[3:4])
  # check - note that data was missing for some explanatory variables especially soil
  cat(paste(focal.var, ":", all.equal(focal.ranges$species, ranges.lookup$species), "\n"))
  ranges.lookup <- cbind(ranges.lookup, focal.ranges[, c(3:4)])
}
```

```
## bio12 : TRUE
## climaticMoistureIndex : TRUE
## monthCountByTemp10 : TRUE
## growingDegDays5 : TRUE
## bio05 : TRUE
## bio06 : TRUE
## bio16 : TRUE
## bio17 : TRUE
```

```r
summary(ranges.lookup)
```

```
##    species                n               bio01_Q05        bio01_Q95
##  Length:22448       Min.   :    20.0   Min.   :-14.91   Min.   :-0.72
##  Class :character   1st Qu.:    35.0   1st Qu.: 15.15   1st Qu.:22.61
##  Mode  :character   Median :    68.0   Median : 18.94   Median :25.95
##                     Mean   :   527.3   Mean   : 18.16   Mean   :24.23
##                     3rd Qu.:   175.0   3rd Qu.: 22.48   3rd Qu.:26.94
##                     Max.   :211181.0   Max.   : 27.76   Max.   :30.46
##    bio12_Q05        bio12_Q95      climaticMoistureIndex_Q05
##  Min.   :   0.0   Min.   :  21.8   Min.   :-1.0000
##  1st Qu.: 733.8   1st Qu.:1647.2   1st Qu.:-0.5100
##  Median :1094.6   Median :2487.5   Median :-0.2400
##  Mean   :1165.8   Mean   :2512.7   Mean   :-0.2314
##  3rd Qu.:1581.2   3rd Qu.:3384.5   3rd Qu.: 0.0600
##  Max.   :5495.6   Max.   :7584.5   Max.   : 0.7300
##  climaticMoistureIndex_Q95 monthCountByTemp10_Q05 monthCountByTemp10_Q95
##  Min.   :-0.9800           Min.   : 0.00          Min.   : 1.90
##  1st Qu.: 0.1800           1st Qu.:10.20          1st Qu.:12.00
##  Median : 0.4600           Median :12.00          Median :12.00
##  Mean   : 0.3326           Mean   :10.46          Mean   :11.78
##  3rd Qu.: 0.5800           3rd Qu.:12.00          3rd Qu.:12.00
##  Max.   : 0.8500           Max.   :12.00          Max.   :12.00
##  growingDegDays5_Q05 growingDegDays5_Q95   bio05_Q05       bio05_Q95
##  Min.   :   0        Min.   : 702.1      Min.   : 7.73   Min.   :14.92
##  1st Qu.:3703        1st Qu.:6429.3      1st Qu.:23.93   1st Qu.:31.34
##  Median :5087        Median :7646.1      Median :26.61   Median :32.92
##  Mean   :4882        Mean   :7027.5      Mean   :26.12   Mean   :32.80
##  3rd Qu.:6379        3rd Qu.:8008.1      3rd Qu.:29.23   3rd Qu.:34.67
##  Max.   :8306        Max.   :9297.1      Max.   :39.88   Max.   :43.51
##    bio06_Q05         bio06_Q95        bio16_Q05        bio16_Q95
##  Min.   :-45.510   Min.   :-24.21   Min.   :   0.0   Min.   :  14.6
##  1st Qu.:  4.600   1st Qu.: 13.54   1st Qu.: 328.0   1st Qu.: 786.9
##  Median :  9.985   Median : 19.01   Median : 481.0   Median :1087.0
##  Mean   :  8.858   Mean   : 16.77   Mean   : 489.5   Mean   :1064.0
##  3rd Qu.: 14.800   3rd Qu.: 21.86   3rd Qu.: 642.1   3rd Qu.:1318.8
##  Max.   : 22.450   Max.   : 23.91   Max.   :2306.5   Max.   :4877.5
##    bio17_Q05        bio17_Q95
##  Min.   :  0.00   Min.   :   0.0
##  1st Qu.: 13.80   1st Qu.: 151.0
##  Median : 41.38   Median : 293.3
##  Mean   : 72.94   Mean   : 343.4
##  3rd Qu.: 95.00   3rd Qu.: 502.1
##  Max.   :932.40   Max.   :1574.0
```

### 4 Custom functions to calculate the number of suitable species

#### 4.1 A first custom function to calculate the number of suitable species

The function has the climatic data of the locations and the filter/ranges data (Q05 and Q95 ranges) of candidate species as inputs, and returns the species richness of suitable species.

```
filter.function <- function(focal.loc, filter.data) {
  filtered.data <- filter.data
  for (f in 1:length(focal.vars)) {
    focal.var <- focal.vars[f]
    LL <- paste0(focal.var, "_Q05")
    filtered.data <- filtered.data[filtered.data[, LL] <= as.numeric(focal.loc[, focal.var]), ]
    UL <- paste0(focal.var, "_Q95")
    filtered.data <- filtered.data[filtered.data[, UL] >= as.numeric(focal.loc[, focal.var]), ]
  }
#  return(filtered.data) # modify the function to return the list of the suitable species
  return(nrow(filtered.data))
}
```

##### 4.2 A second custom function to calculate the number of suitable species in the baseline climate that remain suitable in the future climate

The second filter function sequentially checks the baseline conditions and the future conditions. The result documents the number of species that are suitable first in the baseline and then in the future climate - the species that remain suitable.

```
filter.function2 <- function(base.loc, future.loc, filter.data) {
  filtered.data <- filter.data
  for (f in 1:length(focal.vars)) {
    focal.var <- focal.vars[f]
    LL <- paste0(focal.var, "_Q05")
    filtered.data <- filtered.data[filtered.data[, LL] <= as.numeric(base.loc[, focal.var]), ]
    UL <- paste0(focal.var, "_Q95")
    filtered.data <- filtered.data[filtered.data[, UL] >= as.numeric(base.loc[, focal.var]), ]
  }
  for (f in 1:length(focal.vars)) {
    focal.var <- focal.vars[f]
    LL <- paste0(focal.var, "_Q05")
    filtered.data <- filtered.data[filtered.data[, LL] <= as.numeric(future.loc[, focal.var]), ]
    UL <- paste0(focal.var, "_Q95")
    filtered.data <- filtered.data[filtered.data[, UL] >= as.numeric(future.loc[, focal.var]), ]
  }
#  return(filtered.data) # modify the function to return the list of the suitable species
  return(nrow(filtered.data))
}
```

#### 5 Case Study 1

In the first case study, the subset of environmental ranges (5% and 95% limits) from TreeGOER is used from species where there were a minimum of 20 observations.

##### 5.1 Bioclimatic data for the baseline climate

```
# loc.file <- choose.files()
loc.file <- "C://Data/SuppInfo_001_Tables S1 S2 S4_Data S5 S6.xlsx"
```

```r
loc.data <- data.frame(read_excel(path=loc.file,
                                  sheet="Data S5",
                                  range="A5:Q169"))
# exclude cities with missing data
loc.data <- loc.data[loc.data$INCLUDE == "YES", ]

nrow(loc.data)
```

```
## [1] 161
```

```r
head(loc.data)
```

```
##   ID     ID2      city   country          x          y INCLUDE     bio01 bio12
## 1  1 LOC_001    Aarhus   Denmark  10.186970  56.156817     YES  8.191667   616
## 2  2 LOC_002 Ado Etiki   Nigeria   5.220874   7.626227     YES 25.170834  1343
## 3  3 LOC_003    Albury Australia 146.935862 -36.058745     YES 15.450000   711
## 4  4 LOC_004 Allentown       USA -75.495526  40.599775     YES 10.950000  1133
## 5  5 LOC_005   Amherst       USA -72.516359  42.386430     YES  8.600000  1146
## 6  6 LOC_006  Atakpame      Togo   1.134986   7.531394     YES 26.266666  1298
##   climaticMoistureIndex monthCountByTemp10 growingDegDays5 bio05 bio06 bio16
## 1                  0.04                  5          1626.7  20.8  -3.8   191
## 2                 -0.16                 12          7353.6  32.7  18.8   561
## 3                 -0.45                  9          3801.3  30.8   3.3   230
## 4                  0.05                  6          2748.2  29.9  -7.1   324
## 5                  0.12                  6          2263.4  28.7 -11.2   302
## 6                 -0.24                 12          7760.2  34.5  20.2   524
##   bio17 suitable_baseline
## 1   106               137
## 2    48              2675
## 3   134               195
## 4   239               179
## 5   256               104
## 6    51              1375
```

```r
summary(loc.data)
```

```
##        ID              ID2                 city              country
##  Min.   :  1.00   Length:161         Length:161         Length:161
##  1st Qu.: 42.00   Class :character   Class :character   Class :character
##  Median : 84.00   Mode  :character   Mode  :character   Mode  :character
##  Mean   : 83.26
##  3rd Qu.:124.00
##  Max.   :164.00
##        x                 y               INCLUDE              bio01
##  Min.   :-157.91   Min.   :-43.522   Length:161         Min.   : 3.242
##  1st Qu.: -79.91   1st Qu.:  5.063   Class :character   1st Qu.:10.679
##  Median : -38.53   Median : 32.796   Mode  :character   Median :15.158
##  Mean   : -13.68   Mean   : 21.890                      Mean   :15.694
##  3rd Qu.:  24.99   3rd Qu.: 43.620                      3rd Qu.:20.004
##  Max.   : 172.62   Max.   : 60.216                      Max.   :29.508
##      bio12      climaticMoistureIndex monthCountByTemp10 growingDegDays5
```

```
##  Min.   :  83   Min.   :-0.9500       Min.   : 4.000     Min.   :1348
##  1st Qu.: 657   1st Qu.:-0.3400       1st Qu.: 6.000     1st Qu.:2465
##  Median : 930   Median :-0.0800       Median : 9.000     Median :3696
##  Mean   :1040   Mean   :-0.1278       Mean   : 8.863     Mean   :4149
##  3rd Qu.:1272   3rd Qu.: 0.1000       3rd Qu.:12.000     3rd Qu.:5499
##  Max.   :3422   Max.   : 0.5800       Max.   :12.000     Max.   :8950
##      bio05           bio06             bio16            bio17
##  Min.   :20.40   Min.   :-22.900   Min.   :  44.0   Min.   :  0.0
##  1st Qu.:26.80   1st Qu.: -2.800   1st Qu.: 235.0   1st Qu.: 59.0
##  Median :29.40   Median :  2.600   Median : 324.0   Median :131.0
##  Mean   :29.09   Mean   :  3.201   Mean   : 404.2   Mean   :141.9
##  3rd Qu.:31.50   3rd Qu.:  8.900   3rd Qu.: 509.0   3rd Qu.:188.0
##  Max.   :41.40   Max.   : 23.100   Max.   :1763.0   Max.   :496.0
##  suitable_baseline
##  Min.   :  10.0
##  1st Qu.: 122.0
##  Median : 198.0
##  Mean   : 525.9
##  3rd Qu.: 501.0
##  Max.   :3744.0
```

#### 5.2 Calculate the species richness for the baseline climate with the first filter

The calculations take a few seconds on my laptop.

```
start <- Sys.time()

for (i in 1:nrow(loc.data)) {
# for (i in 1:20) {
  test.result <- filter.function(focal.loc=loc.data[i, ], filter.data=ranges.lookup)
  result <- data.frame(loc.data[i, c("ID2", "suitable_baseline")],
                       suitable_calculated=test.result)
  if (i == 1) {
    out <- result
  }else{
    out <- rbind(out, result)
  }
}

end <- Sys.time()
end - start
```

```
## Time difference of 3.309128 secs
```

We can check now if the newly calculated values correspond to those shown in the article.

```
sum(out$suitable_baseline - out$suitable_calculated)
```

```
## [1] 0
```

#### 5.3 Future bioclimatic data for the cities locations

Load the future climate data.

```
fut.data <- data.frame(read_excel(path=loc.file,
                                  sheet="Data S5",
                                  range="R5:AD169"))

fut.data <- fut.data[fut.data$INCLUDE == "YES", ]

nrow(fut.data)
```

```
## [1] 161
```

```
head(fut.data)
```

```
##        ID INCLUDE bio01 bio12 climaticMoistureIndex monthCountByTemp10
## 1 LOC_001     YES  10.5   639          -0.002589179                  6
## 2 LOC_002     YES  27.2  1455          -0.135550896                 12
## 3 LOC_003     YES  17.4   689          -0.502287772                 11
## 4 LOC_004     YES  13.8  1195           0.010556156                  7
## 5 LOC_005     YES  11.5  1200           0.071638967                  6
## 6 LOC_006     YES  28.6  1324          -0.250673851                 12
##   growingDegDays5 bio05 bio06 bio16 bio17 suitable_filter1 suitable_filter2
## 1         2177.25  23.5  -1.4   200   115              212              104
## 2         8085.15  34.6  20.8   625    49             1059              935
## 3         4526.95  33.2   4.7   219   134              134               91
## 4         3479.50  33.0  -4.1   340   261              129               75
## 5         2931.35  31.5  -7.9   323   276               87               42
## 6         8605.45  36.8  22.5   553    58                3                3
```

```
summary(fut.data)
```

```
##       ID              INCLUDE              bio01           bio12     
##  Length:161         Length:161         Min.   : 5.90   Min.   :  85  
##  Class :character   Class :character   1st Qu.:13.20   1st Qu.: 671  
##  Mode  :character   Mode  :character   Median :17.30   Median : 930  
##                                        Mean   :17.92   Mean   :1050  
##                                        3rd Qu.:22.20   3rd Qu.:1323  
##                                        Max.   :31.60   Max.   :3381  
##  climaticMoistureIndex monthCountByTemp10 growingDegDays5     bio05      
##  Min.   :-0.95304      Min.   : 5.000     Min.   :1898    Min.   :22.60  
##  1st Qu.:-0.38899      1st Qu.: 7.000     1st Qu.:3100    1st Qu.:29.30  
##  Median :-0.13495      Median :10.000     Median :4489    Median :32.10  
##  Mean   :-0.16656      Mean   : 9.503     Mean   :4845    Mean   :31.61  
##  3rd Qu.: 0.05959      3rd Qu.:12.000     3rd Qu.:6289    3rd Qu.:34.20  
##  Max.   : 0.53007      Max.   :12.000     Max.   :9719    Max.   :43.90  
##      bio06             bio16            bio17        suitable_filter1
##  Min.   :-18.500   Min.   :  44.0   Min.   :  0.0   Min.   :   0.0   
##  1st Qu.: -0.100   1st Qu.: 232.0   1st Qu.: 58.0   1st Qu.:  51.0   
##  Median :  4.300   Median : 339.0   Median :132.0   Median : 147.0   
##  Mean   :  5.411   Mean   : 411.2   Mean   :143.3   Mean   : 410.9   
```

```
##  3rd Qu.: 10.800   3rd Qu.: 525.0   3rd Qu.:200.0   3rd Qu.: 310.0
##  Max.   : 24.600   Max.   :1458.0   Max.   :476.0   Max.   :3551.0
##  suitable_filter2
##  Min.   :   0.0
##  1st Qu.:  31.0
##  Median :  85.0
##  Mean   : 258.4
##  3rd Qu.: 163.0
##  Max.   :2976.0
```

#### 5.4 Calculate the species richness for the future data with the first filter

These calculations take less than for the baseline climate, a consequence of narrowing the list of suitable species more quickly.

```
start <- Sys.time()

for (i in 1:nrow(fut.data)) {
# for (i in 1:20) {
  test.result <- filter.function(focal.loc=fut.data[i, ], filter.data=ranges.lookup)
  result <- data.frame(fut.data[i, c("ID", "suitable_filter1")],
                       suitable_calculated=test.result)
  if (i == 1) {
    out <- result
  }else{
    out <- rbind(out, result)
  }
}

end <- Sys.time()
end - start
```

```
## Time difference of 2.499733 secs
```

We can check again now if the newly calculated values correspond to those shown in the article.

```
sum(out$suitable_future - out$suitable_calculated)
```

```
## [1] 0
```

#### 5.5 Calculate the species richness with the second filter

These calculations take just a second or two longer than with the first filter.

```
start <- Sys.time()

for (i in 1:nrow(loc.data)) {
# for (i in 1:20) {
  test.result <- filter.function2(base.loc=loc.data[i, ],
                                  future.loc=fut.data[i, ],
                                  filter.data=ranges.lookup)
```

```
  result <- data.frame(fut.data[i, c("ID", "suitable_filter2")],
                       suitable=test.result)
  if (i == 1) {
    out <- result
  }else{
    out <- rbind(out, result)
  }
}

end <- Sys.time()
end - start
```

```
## Time difference of 2.981174 secs
```

We can check again now if the newly calculated values correspond to those shown in the article.

```
sum(out$suitable_filter2 - out$suitable)
```

```
## [1] 0
```

### 6 Case Study 2

#### 6.1 Baseline bioclimatic data for the 2000 locations

Load the baseline climate data after downloading the supplementary data to a local folder.

```
# loc.file <- choose.files()
loc.file <- "C://Data/SuppInfo_001_Tables S1 S2 S4_Data S5 S6.xlsx"
loc.data <- data.frame(read_excel(path=loc.file,
                                  sheet="Data S6",
                                  range="A5:P2005"))
head(loc.data)
```

```
##   ID      ID2                          Country_GTS         x          y
## 1  1 LOC_0001                                India  84.02083  26.912500
## 2  2 LOC_0002                             Cameroon  12.38750   2.512500
## 3  3 LOC_0003 Congo, The Democratic Republic of the  14.81250  -5.737500
## 4  4 LOC_0004                               Brazil -56.97917  -8.079167
## 5  5 LOC_0005                             Ethiopia  43.60417   8.370833
## 6  6 LOC_0006                               Angola  14.07917 -10.170833
##   country.comment    bio01 bio12 climaticMoistureIndex monthCountByTemp10
## 1            <NA> 24.80417  1372                 -0.15                 12
## 2            <NA> 23.63750  1613                  0.04                 12
## 3            <NA> 24.01250  1290                 -0.16                 12
## 4            <NA> 25.55000  2180                  0.22                 12
## 5            <NA> 22.91250   500                 -0.72                 12
## 6            <NA> 24.58333   623                 -0.57                 12
##   growingDegDays5 bio05 bio06 bio16 bio17 suitable_baseline
## 1          7240.1  37.1   9.0   976    29               154
## 2          6805.2  29.7  18.6   654   135              3920
```

```
## 3          6940.0  30.5  16.0   532     7              1111
## 4          7502.1  34.2  17.8   944    66              2456
## 5          6533.8  31.6  12.9   229    19               269
## 6          7146.1  30.7  16.4   386     1               176
```

```
summary(loc.data) # confirms selection based on Tmo10 and CMI
```

```
##        ID              ID2             Country_GTS              x
##  Min.   :   1.0   Length:2000        Length:2000        Min.   :-174.92
##  1st Qu.: 500.8   Class :character   Class :character   1st Qu.: -53.70
##  Median :1000.5   Mode  :character   Mode  :character   Median :  19.63
##  Mean   :1000.5                                         Mean   :  15.16
##  3rd Qu.:1500.2                                         3rd Qu.:  76.30
##  Max.   :2000.0                                         Max.   : 174.42
##        y             country.comment        bio01           bio12
##  Min.   :-37.112   Length:2000        Min.   :11.48   Min.   : 272.0
##  1st Qu.:-12.981   Class :character   1st Qu.:22.29   1st Qu.: 851.5
##  Median : -1.688   Mode  :character   Median :24.85   Median :1322.0
##  Mean   : -1.321                      Mean   :24.05   Mean   :1468.5
##  3rd Qu.: 10.012                      3rd Qu.:26.27   3rd Qu.:1879.2
##  Max.   : 42.979                      Max.   :29.58   Max.   :7199.0
##  climaticMoistureIndex monthCountByTemp10 growingDegDays5     bio05
##  Min.   :-0.7900       Min.   :12         Min.   :2366    Min.   :18.30
##  1st Qu.:-0.5100       1st Qu.:12         1st Qu.:6304    1st Qu.:30.70
##  Median :-0.1700       Median :12         Median :7245    Median :32.30
##  Mean   :-0.1564       Mean   :12         Mean   :6953    Mean   :32.77
##  3rd Qu.: 0.1525       3rd Qu.:12         3rd Qu.:7766    3rd Qu.:34.70
##  Max.   : 0.7900       Max.   :12         Max.   :8973    Max.   :44.50
##      bio06           bio16            bio17         suitable_baseline
##  Min.   : 1.10   Min.   :  81.0   Min.   :   0.0   Min.   :   0.0
##  1st Qu.:10.80   1st Qu.: 469.0   1st Qu.:  13.0   1st Qu.: 188.0
##  Median :15.40   Median : 642.5   Median :  42.0   Median : 909.5
##  Mean   :14.68   Mean   : 676.0   Mean   : 115.9   Mean   :1452.4
##  3rd Qu.:18.90   3rd Qu.: 854.0   3rd Qu.: 150.5   3rd Qu.:2536.5
##  Max.   :23.60   Max.   :3326.0   Max.   :1511.0   Max.   :5252.0
```

#### 6.2 Calculate the species richness for the baseline climate

For the second case study, only the first filter is used (but there are no practical reasons why the second filter could not be used). The calculations take around a minute on my laptop.

```
start <- Sys.time()

for (i in 1:nrow(loc.data)) {
# for (i in 1:20) {
  test.result <- filter.function(focal.loc=loc.data[i, ], filter.data=ranges.lookup)
  result <- data.frame(loc.data[i, c("ID2", "suitable_baseline")],
                       suitable_calculated=test.result)
  if (i == 1) {
    out <- result
  }else{
    out <- rbind(out, result)
```

```
  }
}

end <- Sys.time()
end - start
```

```
## Time difference of 55.64457 secs
```

We can check now if the newly calculated values correspond to those shown in the article.

```
sum(out$suitable_baseline - out$suitable_calculated)
```

```
## [1] 0
```

##### 6.3 Future bioclimatic data for the 2000 locations

Load the future climate data.

```
loc.data <- data.frame(read_excel(path=loc.file,
                                  sheet="Data S6",
                                  range="S5:AD2005"))
head(loc.data)
```

```
##         ID bio01 bio12 climaticMoistureIndex monthCountByTemp10 growingDegDays5
## 1 LOC_0001  26.2  1494           -0.07455441                 12         7757.90
## 2 LOC_0002  25.6  1673            0.05081011                 12         7522.00
## 3 LOC_0003  26.1  1346           -0.17998348                 12         7709.40
## 4 LOC_0004  27.8  2044            0.11111098                 12         8311.65
## 5 LOC_0005  25.1   528           -0.72038839                 12         7330.25
## 6 LOC_0006  26.5   678           -0.54844379                 12         7839.35
##   bio05 bio06 bio16 bio17 zone_2050s suitable_future
## 1  38.2  10.6  1073    28      humid             104
## 2  31.8  20.5   691   148      humid            3431
## 3  32.6  18.2   554     8      humid            1125
## 4  36.7  20.3   916    61      humid             106
## 5  33.5  15.6   232    21        dry             209
## 6  32.5  18.3   420     1        dry             113
```

```
summary(loc.data)
```

```
##       ID                 bio01           bio12        climaticMoistureIndex
##  Length:2000        Min.   :12.60   Min.   : 262.0   Min.   :-0.8122
##  Class :character   1st Qu.:24.40   1st Qu.: 888.5   1st Qu.:-0.5081
##  Mode  :character   Median :26.80   Median :1355.5   Median :-0.2028
##                     Mean   :26.06   Mean   :1488.1   Mean   :-0.1765
##                     3rd Qu.:28.30   3rd Qu.:1896.2   3rd Qu.: 0.1250
##                     Max.   :31.80   Max.   :7269.0   Max.   : 0.7887
##  monthCountByTemp10 growingDegDays5     bio05           bio06
##  Min.   :12         Min.   :2778    Min.   :18.40   Min.   : 2.90
##  1st Qu.:12         1st Qu.:7087    1st Qu.:32.80   1st Qu.:12.90
```

```
##  Median :12         Median :7965    Median :34.60   Median :17.60
##  Mean   :12         Mean   :7686    Mean   :34.98   Mean   :16.71
##  3rd Qu.:12         3rd Qu.:8489    3rd Qu.:37.00   3rd Qu.:21.10
##  Max.   :12         Max.   :9767    Max.   :48.30   Max.   :25.20
##      bio16             bio17          zone_2050s        suitable_future
##  Min.   :  77.0   Min.   :   0.00   Length:2000        Min.   :   0.00
##  1st Qu.: 495.0   1st Qu.:  12.75   Class :character   1st Qu.:  10.75
##  Median : 688.0   Median :  42.00   Mode  :character   Median : 152.50
##  Mean   : 703.4   Mean   : 113.10                      Mean   : 669.71
##  3rd Qu.: 877.2   3rd Qu.: 151.25                      3rd Qu.: 937.75
##  Max.   :3226.0   Max.   :1518.00                      Max.   :4904.00
```

#### 6.4 Calculate the species richness for the future data

These calculations take less than for the baseline climate, a consequence of narrowing the list of suitable species more quickly.

```
start <- Sys.time()

for (i in 1:nrow(loc.data)) {
# for (i in 1:20) {
  test.result <- filter.function(focal.loc=loc.data[i, ], filter.data=ranges.lookup)
  result <- data.frame(loc.data[i, c("ID", "suitable_future")],
                       suitable_calculated=test.result)
  if (i == 1) {
    out <- result
  }else{
    out <- rbind(out, result)
  }
}

end <- Sys.time()
end - start
```

```
## Time difference of 35.58618 secs
```

We can check again now if the newly calculated values correspond to those shown in the article.

```
sum(out$suitable_future - out$suitable_calculated)
```

```
## [1] 0
```

### 7 Session information

```
sessionInfo()
```

```
## R version 4.2.1 (2022-06-23 ucrt)
## Platform: x86_64-w64-mingw32/x64 (64-bit)
## Running under: Windows 10 x64 (build 19045)
```

```
##
## Matrix products: default
##
## locale:
## [1] LC_COLLATE=English_United Kingdom.utf8
## [2] LC_CTYPE=English_United Kingdom.utf8
## [3] LC_MONETARY=English_United Kingdom.utf8
## [4] LC_NUMERIC=C
## [5] LC_TIME=English_United Kingdom.utf8
##
## attached base packages:
## [1] stats     graphics  grDevices utils     datasets  methods   base
##
## other attached packages:
## [1] readxl_1.4.1      data.table_1.14.2
##
## loaded via a namespace (and not attached):
##  [1] fansi_1.0.3      utf8_1.2.2       digest_0.6.29    cellranger_1.1.0
##  [5] lifecycle_1.0.3  magrittr_2.0.3   evaluate_0.16    pillar_1.8.1
##  [9] rlang_1.0.6      stringi_1.7.8    cli_3.4.1        rematch_1.0.1
## [13] rstudioapi_0.14  vctrs_0.5.1      rmarkdown_2.16   tools_4.2.1
## [17] stringr_1.4.1    glue_1.6.2       xfun_0.33        yaml_2.3.5
## [21] fastmap_1.1.0    compiler_4.2.1   pkgconfig_2.0.3  htmltools_0.5.3
## [25] knitr_1.40       tibble_3.1.8
```
